# Comprehensive Characterization and Global Transcriptome Analyses of Human Fetal Liver Terminal Erythropoiesis

**DOI:** 10.1101/2023.06.15.545026

**Authors:** Yongshuai Han, Shihui Wang, Yaomei Wang, Yumin Huang, Chengjie Gao, Xinhua Guo, Lixiang Chen, Huizhi Zhao, Xiuli An

## Abstract

The fetal liver (FL) is the key erythropoietic organ during fetal development, but knowledge on human FL erythropoiesis is very limited. In this study, we sorted primary erythroblasts from FL cells and performed RNA sequencing analyses. We found that temporal gene expression patterns reflected changes in function during primary human FL terminal erythropoiesis. Notably, expression of genes enriched in proteolysis and autophagy was upregulated in orthochromatic erythroblasts (OrthoE), suggesting involvement of these pathways in enucleation. We also performed RNA sequencing of *in vitro* cultured erythroblasts derived from FL CD34^+^ cells. Comparison of transcriptomes between the primary and cultured erythroblasts revealed significant differences, indicating impacts of the culture system on gene expression. Notably, lipid metabolism gene expression was increased in cultured erythroblasts. We further immortalized erythroid cell lines from FL and cord blood (CB) CD34^+^ cells (FL-iEry and CB-iEry, respectively). FL-iEry and CB-iEry are immortalized at the proerythroblast stage and can be induced to differentiate into OrthoE, but their enucleation ability is very low. Comparison of transcriptomes between OrthoE with and without enucleation capability revealed downregulation of pathways involved in chromatin organization and mitophagy in OrthoE without enucleation capacity, indicating that defects in chromatin organization and mitophagy contribute to the inability of OrthoE to enucleate. Additionally, the expression levels of *HBE1*, *HBZ*, and *HBG2* were upregulated in FL-iEry compared with CB-iEry, and this was accompanied by downregulation of *BCL11A* and upregulation of *LIN28B* and *IGF2BP1*. Our study provides new insights into human FL erythropoiesis and rich resources for future studies.

## Introduction

Erythropoiesis is a process by which red blood cells are produced. It occurs first in the yolk sac as the “primitive” form, then is gradually replaced by the “definitive” form in the fetal liver (FL) and bone marrow (BM) during fetal development and after birth [1, 2]. The earliest committed erythroid progenitor is the burst forming unit-erythroid (BFU-E); the progenitors differentiate into a late-stage erythroid progenitor, the colony-forming unit-erythroid (CFU-E). The CFU-E undergoes terminal erythroid differentiation to become erythroid precursors including proerythroblasts (ProE), basophilic erythroblasts (BasoE), polychromatic erythroblasts (PolyE), and orthochromatic erythroblasts (OrthoE) [3, 4]. During terminal erythroid differentiation, cell size progressively decreases while chromatin condensation and hemoglobin synthesis increase. Finally, OrthoE expel their nuclei to become reticulocytes, which mature into red blood cells in the blood stream [5]. These changes enable morphologic differentiation of erythroblasts at different developmental stages.

For more than 10 years, flow cytometry-based methods using surface markers to separate both mouse and human erythroid cells at each developmental stage have been developed by us and others [3, 6–10]. These novel methods have enabled the study of normal and disordered erythropoiesis in a stage-specific manner [11–16]. Global transcriptomic, proteomic, and epigenetic analyses of these cell populations have provided novel insights into the biology of erythropoiesis [17–19]. Our previous transcriptome analyses of human and murine erythroblasts revealed significant stage- and species-specific differences across stages of terminal erythroid differentiation [17]. The epigenic landscape of human erythropoiesis reveals that erythroid cells exhibit chromatin accessibility patterns distinct from other cell types. It also reveals stage-specific patterns of regulation [19]. However, it should be noted that for human erythropoiesis, all the omics analyses were performed on *in vitro* cultured erythroid cells derived from cord blood (CB) or peripheral blood (PB) CD34^+^ cells. In marked contrast, there is a lack of knowledge on human FL erythropoiesis.

In the present study, we first showed that the surface markers we identified for separating cultured human erythroblasts [3] can be used to sort out primary erythroblasts at different developmental stages from human FL cells. We then differentiated human FL CD34^+^ cells to erythroid cells *in vitro* using a three-phase erythroid culture system and showed that the *in vitro* erythropoiesis profile of FL CD34^+^ cells is more similar to that of CB CD34^+^ cells than to that of PB CD34^+^ cells. We performed RNA sequencing (RNA-seq) analyses on erythroblasts at each developmental stage sorted from both human FL and *in vitro* cultured erythroblasts derived from human FL CD34^+^ cells. Comparison of the transcriptome between human primary FL erythroblasts and cultured erythroblasts derived from human FL CD34^+^ cells revealed many differences in gene expression between the primary human FL erythroblasts and cultured erythroblasts, indicating the influence of the culture system on terminal erythropoiesis. We also established immortalized erythroid cell lines from human FL cells and CB CD34^+^ cells, which we termed FL-iEry and CB-iEry, respectively. Characterization and RNA-seq analyses revealed that both FL-iEry and CB-iEry were immortalized at the ProE stage and could be induced to undergo terminal erythroid differentiation. Interestingly, the expression levels of embryonic and γ-globin genes were upregulated in FL-iEry compared with CB-iEry. Together, our findings provide novel insights into human FL erythropoiesis. Establishment of an FL-iEry cell line should facilitate studies in human FL erythropoiesis.

## Results

### Combination of surface markers enables separation of primary erythroblasts from human FL cells

Using a three-phase erythroid culture system, we developed a flow cytometry-based method for separating human erythroblasts at each distinct developmental stage using glycophorin A (GPA), band 3, and α4 integrin as surface markers [3, 20–22]. This method can be used to separate primary erythroblasts at different stages of terminal erythroid differentiation in the human BM with high purity [3]. We examined whether the combination of these markers can separate primary erythroblasts in the human FL. Figure S1A shows the representative expression profile of band 3 versus α4 integrin of the GPA^+^ cells. Based on the expression levels of band 3 and α4 integrin, five clusters were gated and sorted. Cytospin analyses of the sorted cells showed that they morphologically resembled ProE, early BasoE, late BasoE, PolyE, and OrthoE (Figure S1B). These results demonstrate that the combination of the surface markers GPA, band 3, and α4 integrin can be used to obtain primary human FL erythroblasts at each distinct stage using fluorescence-activated cell sorting (FACS).

### Overall transcriptome profiles of primary human FL erythroblasts confirm FACS based separation

To examine the changes in gene expression patterns during terminal differentiation of human FL cells *in vivo*, we obtained the transcriptome of the sorted primary human FL erythroblasts of each stage across terminal erythropoiesis. Consistent with our previous report [17], decreasing numbers of expressed genes were detected from ProE to OrthoE with 10,215, 9060, 7808, and 5750 expressed genes (≥1 transcript per million in at least one stage). All the expressed genes are listed in Table S1. Principal component analysis (PCA) showed clear separation of samples from each stage with the exception of one sample from PolyE (**Figure 1**A). Distance analysis revealed a shorter distance between samples within each stage and a longer distance between samples from different stages (Figure 1B). Pairwise correlation analysis revealed decreasing similarity in gene expression profiles during the differentiation process, from 0.95 between ProE and BasoE to 0.82 between ProE and OrthoE (Figure 1C).

**Figure 1.**
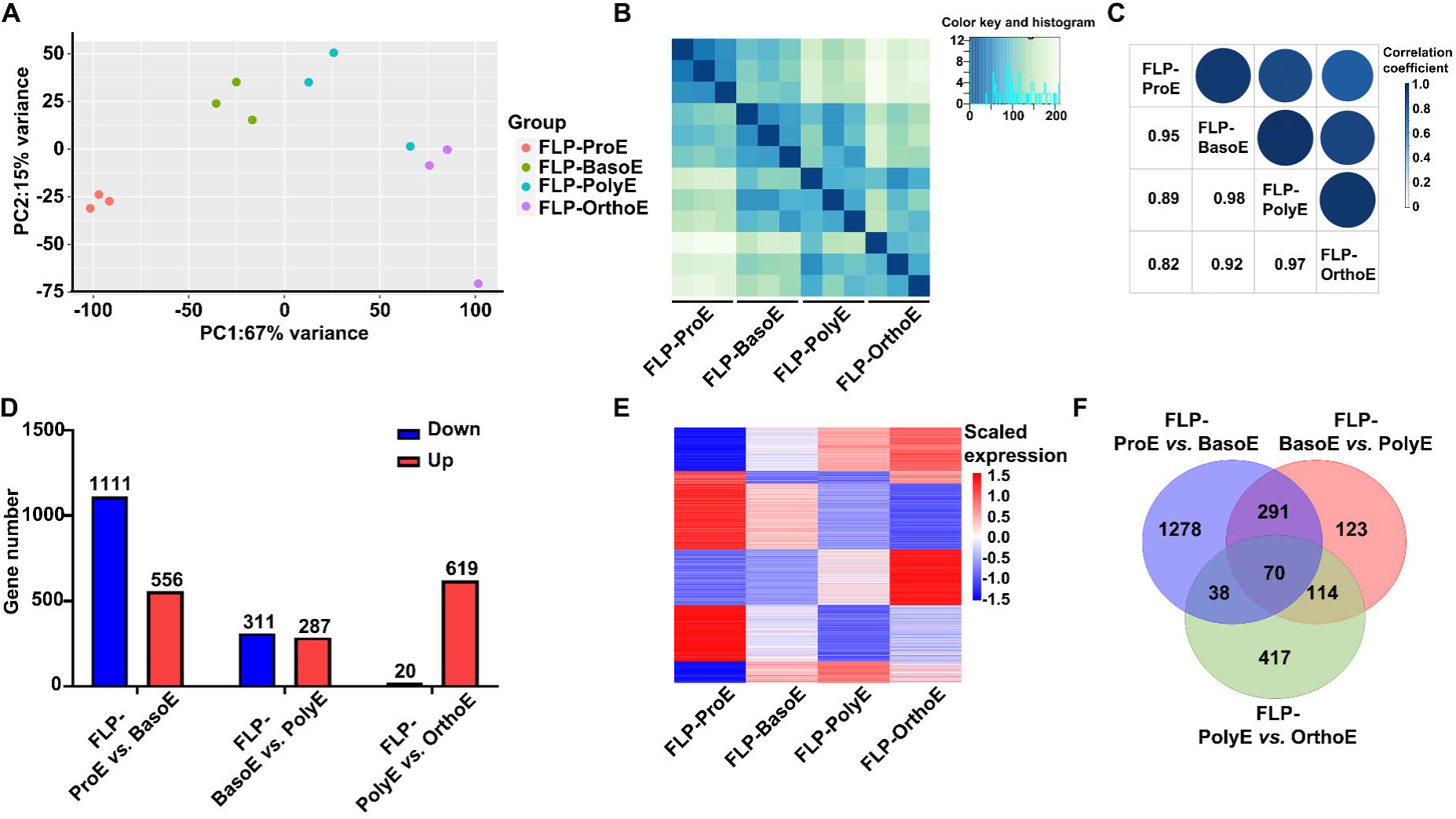
RNA-seq analyses of primary erythroblasts from human fetal liver. **A.** Principal component analyses showing the separation of erythroblasts at distinct developmental stage from fetal liver. **B.** Heat map of pair-wise distances between samples. **C.** Mixed plot of pair-wise correlation coefficients between stages of human fetal liver primary erythroblasts. **D.** Bar plot of DEGs numbers between adjacent stages. **E.** Heat map representation of gene expression of all DEGs from adjacent stage. **F.** Venn diagram of DEGs from adjacent stage. DEGs, differentially expressed genes; FLP-ProE, fetal liver primary proerythroblasts; FLP-BasoE, fetal liver primary basophilic erythroblasts; FLP-PolyE, fetal liver primary polychromatic erythroblasts; FLP-OrthoE, fetal liver primary orthochromatic erythroblasts.

We then compared gene expression between adjacent stages and obtained a total of 2331 differentially expressed genes (DEGs), with 1667 DEGs between ProE and BasoE, 598 DEGs between BasoE and PolyE, and 639 DEGs between PolyE and OrthoE (Figure 1D). A heat map representation of all DEGs is shown in Figure 1E. A list of DEGs between adjacent stages is provided in Table S2. Notably, the greatest changes in gene expression (almost 70% of all DEGs) were seen between ProE and BasoE, indicating that the most notable changes occur in the early stage of human FL terminal erythropoiesis *in vivo*. A Venn diagram of DEGs between adjacent stages (ProE *vs* BasoE, BasoE *vs* PolyE, and PolyE *vs* OrthoE) showed that 77% of DEGs identified in ProE versus BasoE were only found in one comparison (Figure 1F), suggesting stage-specific changes from ProE to BasoE.

### Temporal patterns of gene expression during primary human FL terminal erythropoiesis reflect changes in function

Dramatic changes occur during terminal erythropoiesis. To examine the dynamic changes in gene expression during this process, we performed a time-course analysis of the gene expression of all 2331 DEGs identified from the pairwise comparison of adjacent stages, followed by gene ontology (GO) and pathway analyses of the identified clusters. Heat maps representing the gene expression patterns and their enriched GO terms are shown in **Figure 2**. The expression of genes in cluster 1 increased from ProE to OrthoE, with significant upregulation in OrthoE. These genes are involved in proteolysis and autophagy, suggesting increasing need for clearance of protein and organelles before enucleation. The genes in cluster 2, which showed decreasing expression from ProE to OrthoE, were enriched in the non-coding RNA and transfer RNA metabolic process, indicating a decrease in protein synthesis. The genes in cluster 3, which showed progressively increasing expression from ProE to OrthoE, were enriched in histone deacetylases, porphyrin-containing compound metabolic processes, and erythrocyte differentiation, consistent with increased hemoglobin synthesis and chromatin remodeling during terminal erythropoiesis. Gene expression in cluster 4 was characterized by abundant expression in ProE and decreasing expression in later stages. The genes in this group were enriched in positive regulation of cell adhesion, consistent with previous findings that adhesion molecules are decreased during human terminal erythropoiesis [3]. The expression of genes in cluster 5 was upregulated from ProE to BasoE, then remained stable from BasoE to OrthoE. The expression of these genes is enriched in the cell cycle, implying more active cell cycle in ProE than in later stages.

**Figure 2.**
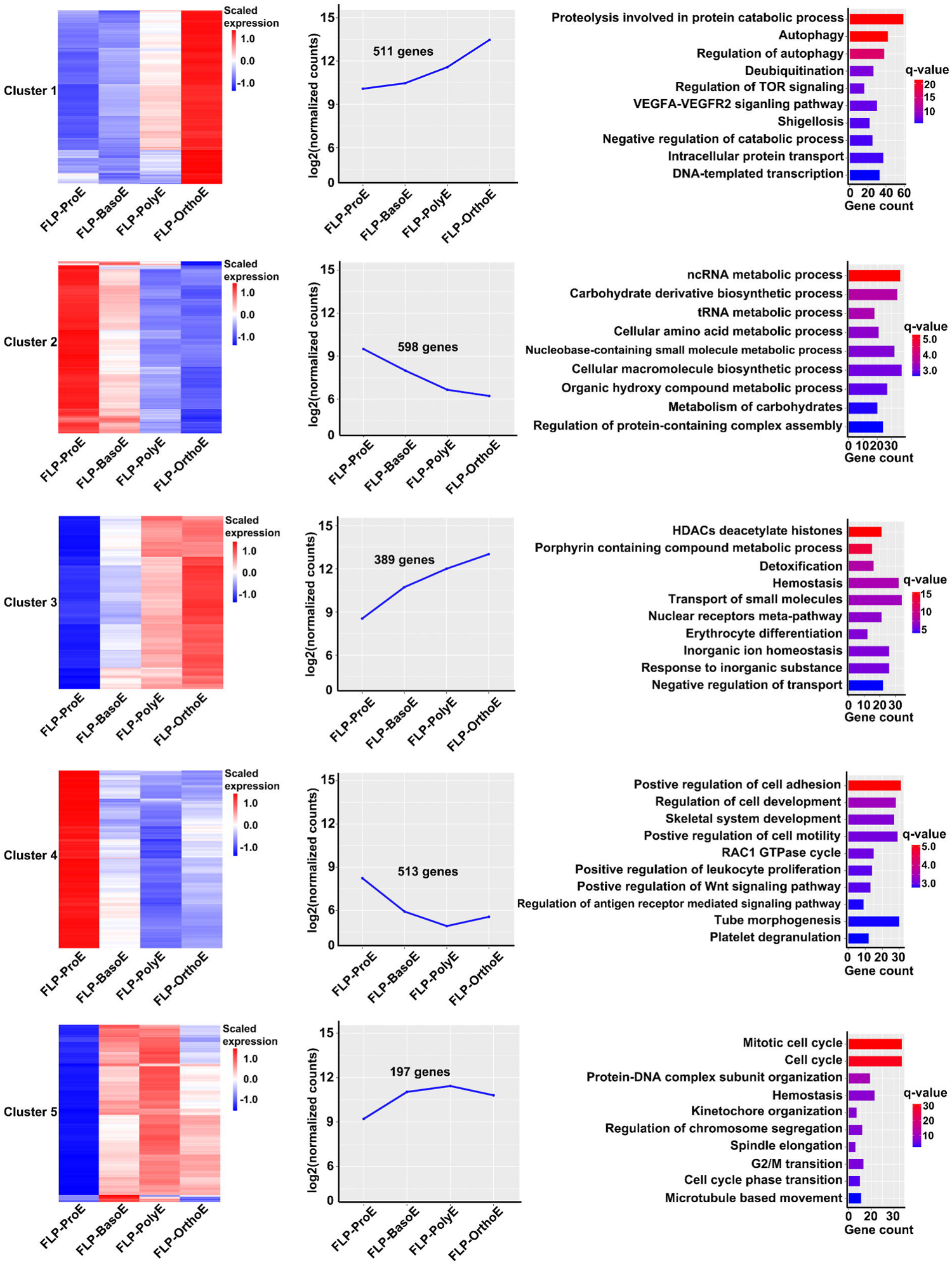
Clusters of differentially expressed genes across stages during primary human fetal liver terminal erythropoiesis. The gene expression of five identified clusters of all DEGs from adjacent stage comparison are shown in a heat map (left panel) and curve plot (middle panel). Enriched GO and pathway terms of genes within clusters are shown in a bar plot (right panel). The q-value represents the log-transformed adjusted *P*.

### *In vitro* erythropoiesis profile of human FL CD34^+^ cells is more similar to that of CB CD34^+^ cells than that of PB CD34^+^ cells

Many previous studies used CB-, PB-, or BM-derived CD34^+^ cells to model neonatal or adult human erythropoiesis [23–25]. In the present study, we differentiated human FL CD34^+^ cells to the erythroid lineage using the three-phase erythroid culture system described in our previous studies [3, 20–22]. **Figure 3**A shows the growth curve of FL CD34^+^ cell-derived erythroid cells, showing that the cell population expanded about 263,000-fold over the 15-day culture period. We previously reported that BFU-E and CFU-E cells are immunophenotypically defined as GPA^−^IL-3R^−^CD34^+^CD36^−^ and GPA^−^IL-3R^−^CD34^−^CD36^+^, respectively [8]. Using this flow cytometry-based strategy, we monitored early-stage erythropoiesis during the first 7 days of culture. The representative expression profiles of GPA versus IL-3R are shown in the upper panel of Figure 3B, which reveals gradual acquisition of GPA accompanied by a decrease in IL-3R expression. The expression profiles of CD34 versus CD36 are shown in the lower panel of Figure 3B, which reveals gradual loss of CD34 expression accompanied by gradual gain of CD36 expression. Yan et al. showed that a subset of CD34^+^CD36^+^ cells, phenotypically located in the transition between BFU-E and CFU-E, were present with notable abundance in PB-but not CB-derived cultures from days 3 to 6 of culture [10]. Notably, FL-derived cultures presented a pattern similar to that of CB-derived cultures, indicating close similarity between FL and CB. We also examined the erythroid colony-forming ability of erythroid progenitors. Figure 3C shows that while BFU-E colonies peaked on day 3, CFU-E colonies peaked on day 5.

**Figure 3.**
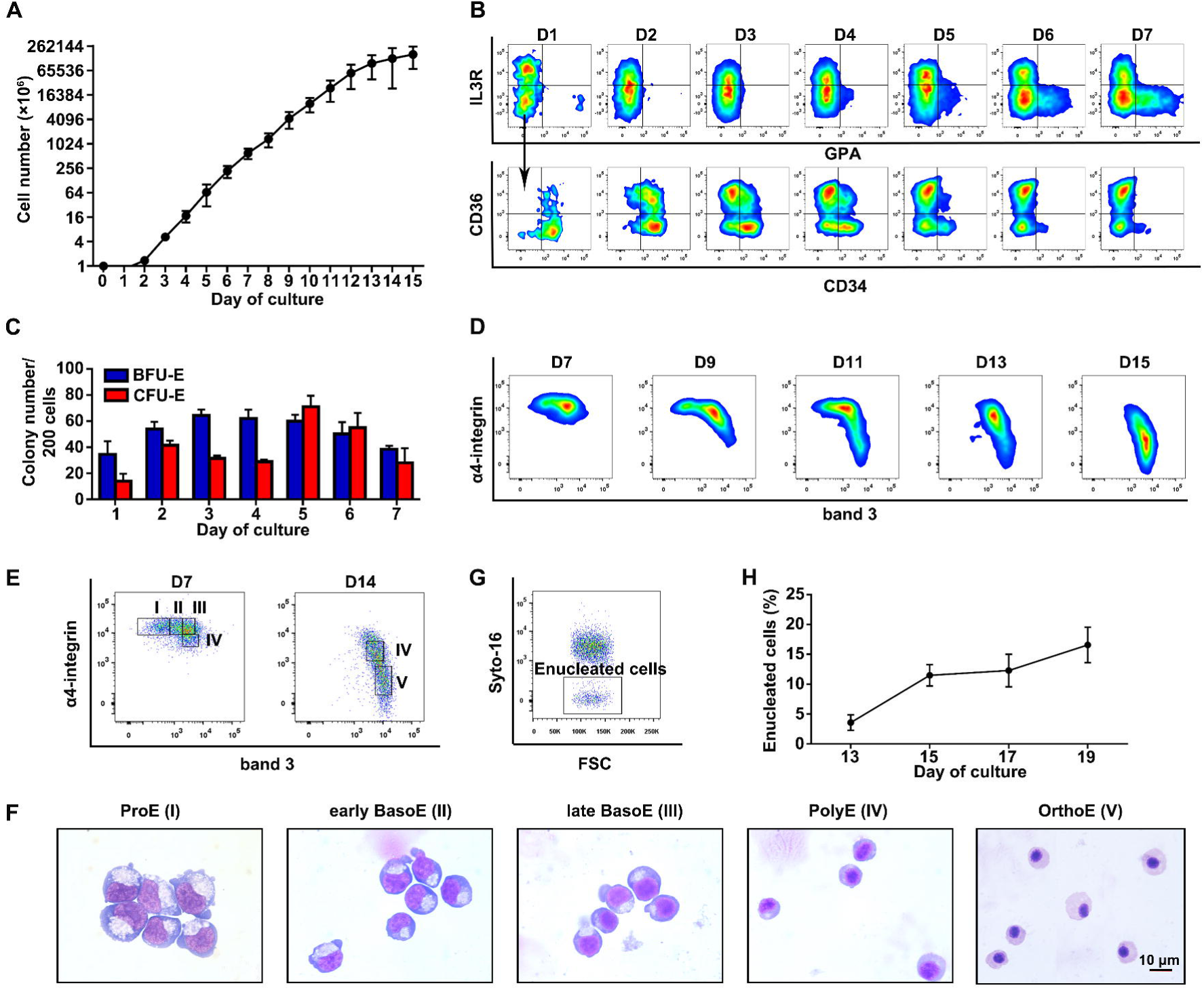
*In vitro* erythropoiesis profile of human fetal liver CD34^+^ cells. **A.** Growth curve of the FL CD34^+^ cell-derived erythroid cells. Error bars indicate standard deviation (n = 6). **B.** Representative dot plots of GPA versus IL-3R (upper panel) and CD34 versus CD36 of GPA^-^IL-3R^-^ cells from cultured day 1 to day 7 (lower panel). **C.** Quantitation of colony forming ability of BFU-E and CFU-E on indicated days. Error bars indicate standard deviation (n = 3). **D.** Representative dot plots of band 3 versus α4-integrin of GPA^+^ cells from cultured day 7 to day 15. **E.** Gating strategy of terminal erythroblasts on cultured day 7 or day 14 based on the expression levels of band 3 and α4-integrin of all GPA^+^ cells. Population I (ProE) was band3^neg^ α4-integrin^hi^, population II (early BasoE) was band3^low^ α4-integrin^hi^, population III (late BasoE) was band3^med^ α4-integrin^hi^, population IV (PolyE) was band3^med^ α4-integrin^med^, and population V (OrthoE) was band3^hi^ α4-integrin^low^. **F.** Representative images of sorted erythroblasts from above five populations shown in Figure 3E. Scale bar = 10µm. **G.** Representative enucleation profile on cultured day 17. **H.** Quantitative analyses of enucleation. Error bars indicate standard deviation (n = 3).

Next, we examined the terminal erythroid differentiation of FL CD34^+^ cells by flow cytometry using GPA, band 3, and α4 integrin as surface markers [3, 20, 26]. The expression profiles of band 3 versus α4 integrin of GPA^+^ cells are shown in Figure 3D. As terminal erythroid differentiation proceeded, band 3 expression increased while α4 integrin expression decreased. Based on the expression levels of band 3 and α4 integrin, we sorted ProE, early BasoE, and late BasoE from cells cultured for 7 days and sorted PolyE and OrthoE from cells cultured for 14 days. The gating strategies are shown in Figure 3E. Cytospin analyses of the sorted cells showed that the sorted cells morphologically resembled ProE, early BasoE, late BasoE, PolyE, and OrthoE, respectively (Figure 3F). These results indicate that the combination of GPA, band 3, and α4-integrin can be used to monitor terminal differentiation of FL CD34^+^ cells and isolate erythroblasts derived from FL CD34^+^ cells.

Enucleation is the last step of terminal erythroid differentiation. Therefore, we assessed enucleation using SYTO 16 staining by flow cytometry. The representative enucleation profiles and quantitative analyses are shown in Figure 3G and H, respectively. Unexpectedly, on day 17, the enucleation rate of erythroblasts derived from FL CD34^+^ cells only reached <20% which is significantly lower than that of erythroblasts derived from PB, CB, or BM CD34^+^ cells (the enucleation rate of which ranged from 45% to 65%) [20, 26, 27].

### Overall transcriptome profiles of cultured FL-derived erythroblasts reveal many more DEGs during *in vitro* terminal erythropoiesis

Next, we performed RNA-seq analyses of the cultured erythroblasts derived from human FL CD34^+^ cells. On average, the numbers of expressed genes were 10,115, 9620, 7240, and 5819 in the cultured ProE, BasoE, PolyE, and OrthoE, respectively, and these numbers were comparable to those of primary erythroblasts at the same stage. A list of expressed genes is provided in Table S3. PCA showed clear separation of the four stages (**Figure 4**A). We also analyzed the transcriptome profiles of primary and cultured erythroblasts using all expressed genes together and found different trajectories of *in vivo* and *in vitro* terminal erythropoiesis (Figure S2). The distance analysis showed that different from the continuous separation of each stage of primary erythroblasts (see Figure 1B), the cultured erythroblasts exhibited two-part separation, with ProE/BasoE as one part and PolyE/OrthoE as the other part (Figure 4B). Pairwise correlation analysis showed that the correlation coefficient between the cultured BasoE and PolyE was the smallest among all correlation coefficients between adjacent stages (Figure 4C), indicating more changes from BasoE and PolyE in the culture system. Pairwise transcriptome comparison between adjacent stages in cultured erythropoiesis revealed 5263 DEGs in total (Table S4), which was a much larger number than that in primary erythroblasts (2331 DEGs). Notably, of the total DEGs, 77% (4058 of 5263) were identified between the cultured BasoE and PolyE, whereas only 259 DEGs were identified between the cultured ProE and BasoE (Figure 4D). This indicated high similarity between the cultured ProE and BasoE but dramatic differences between the cultured BasoE and PolyE. A heat map representing gene expression of all DEGs is shown in Figure 4E. Venn diagram analyses revealed that 83% of DEGs in cultured erythroblasts were only identified in one comparison (Figure 4F), indicating strong stage-specific changes from BasoE to PolyE. The many differences described above suggest different differentiation patterns of primary and cultured terminal erythropoiesis, most likely due to the effects of the *in vitro* culture environment.

**Figure 4.**
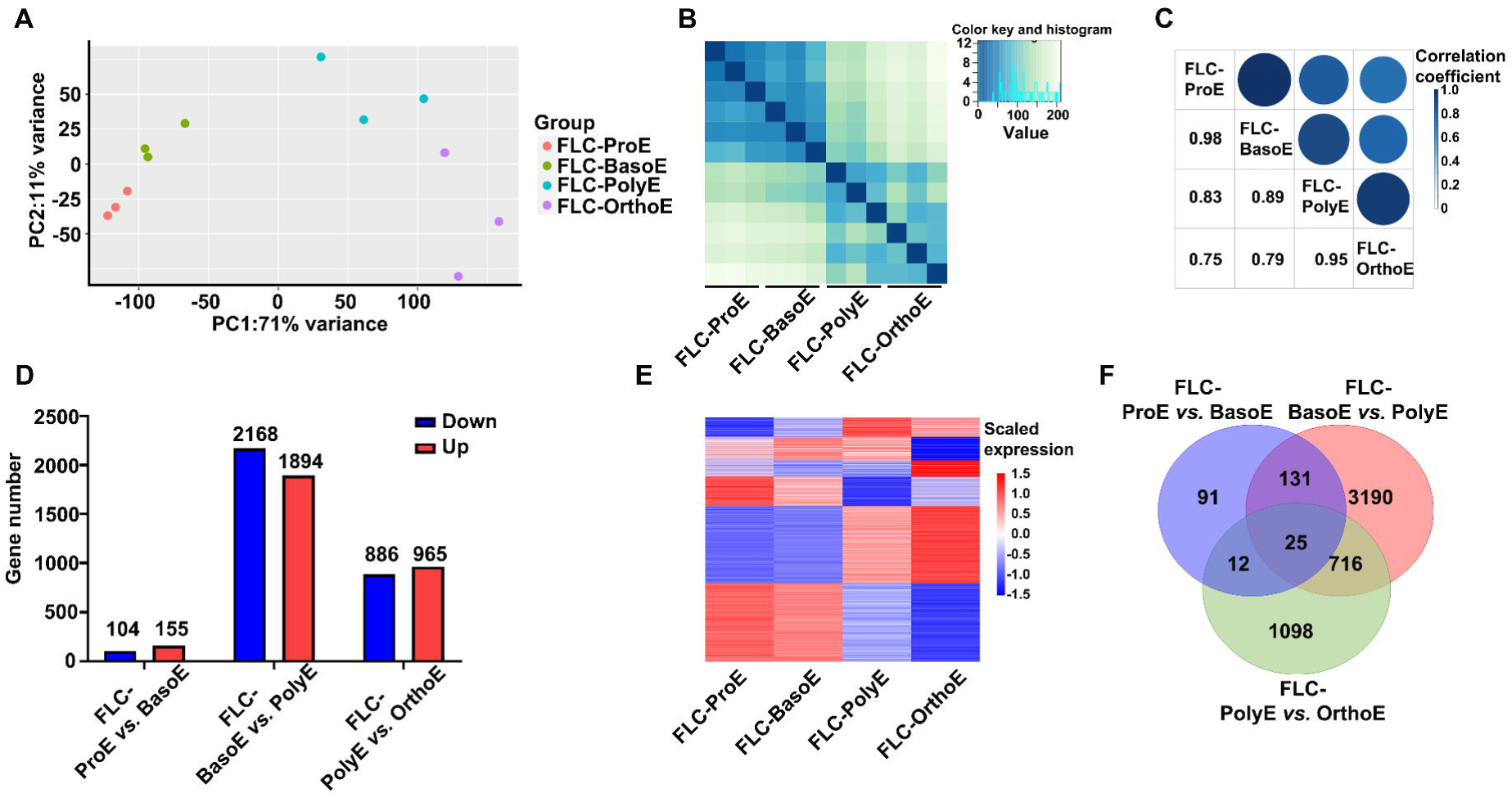
RNA-seq analyses of cultured erythroblasts from human fetal liver CD34^+^ cells. **A.** Principal component analyses showing the separation of fetal liver cultured erythroblasts. **B.** Heat map of pair-wise distances between each stage of terminal erythroblasts. **C.** Mixed plot of pair-wise correlation coefficients between stages. **D.** Bar plot of DEG numbers of adjacent stages. **E.** Heat map representation of gene expression of all DEGs from adjacent stage comparison. **F.** Venn diagram of DEGs from adjacent stages. FLC-ProE, fetal liver cultured proerythroblasts; FLC-BasoE, fetal liver cultured basophilic erythroblasts; FLC-PolyE, fetal liver cultured polychromatic erythroblasts; FLC-OrthoE, fetal liver cultured orthochromatic erythroblasts.

### Predominant changes at PolyE and OrthoE stages during *in vitro* terminal erythropoiesis

We next performed a time-course analysis of the expression patterns of all 5263 DEGs identified above to examine the dynamic patterns of gene expression during the *in vitro* terminal erythropoiesis of human FL CD34^+^ cells. As shown in **Figure 5**, six clusters were identified. Of these six clusters, only two were similar to the clusters of terminal erythropoiesis of primary human FL cells. In cluster 1, expression of genes increased from ProE to OrthoE, and these genes were enriched in proteolysis and autophagy (similar to cluster 1 in primary erythropoiesis). In cluster 2, expression of genes decreased from ProE to OrthoE, and these genes were enriched in the non-coding RNA, DNA, and transfer RNA metabolic process (similar to gene cluster 2 in primary erythropoiesis). In cluster 3, the expression of genes increased from ProE to PolyE but slightly decreased in OrthoE, and these genes were enriched in hemostasis, erythrocyte differentiation, and phospholipid metabolism. In cluster 4, expression of genes decreased from ProE to PolyE but slightly increased in OrthoE, and these genes were enriched in mitochondrial gene expression, mitochondrial RNA metabolic processes, and mitochondrion organization. In cluster 5, expression of genes was relatively stable from ProE to PolyE and specifically upregulated in OrthoE, and these genes were enriched in the cellular response to topologically incorrect protein, regulation of the apoptotic signaling pathway, and mitochondrion organization. Genes in cluster 6 were decreased and specifically downregulated in OrthoE but enriched in portions of the cell cycle such as G2/M transition and DNA metabolic processes. The patterns and GO enrichment of clusters 3 to 6 were not noted in human FL primary erythropoiesis but were similar to those reported in terminal erythropoiesis of CB CD34^+^ cells [17], demonstrating similarity of erythroblasts cultured from CB CD34^+^ and FL CD34^+^ cells. The stage-specific upregulated genes in each stage in FL *in vitro* and *in vivo* terminal erythropoiesis are listed in Table S5. Together, these results demonstrate that although there are conserved changes during *in vivo* and *in vitro* terminal erythropoiesis of human FL, *in vitro* culture leads to specific changes, particularly in the very late stage of terminal erythropoiesis.

**Figure 5.**
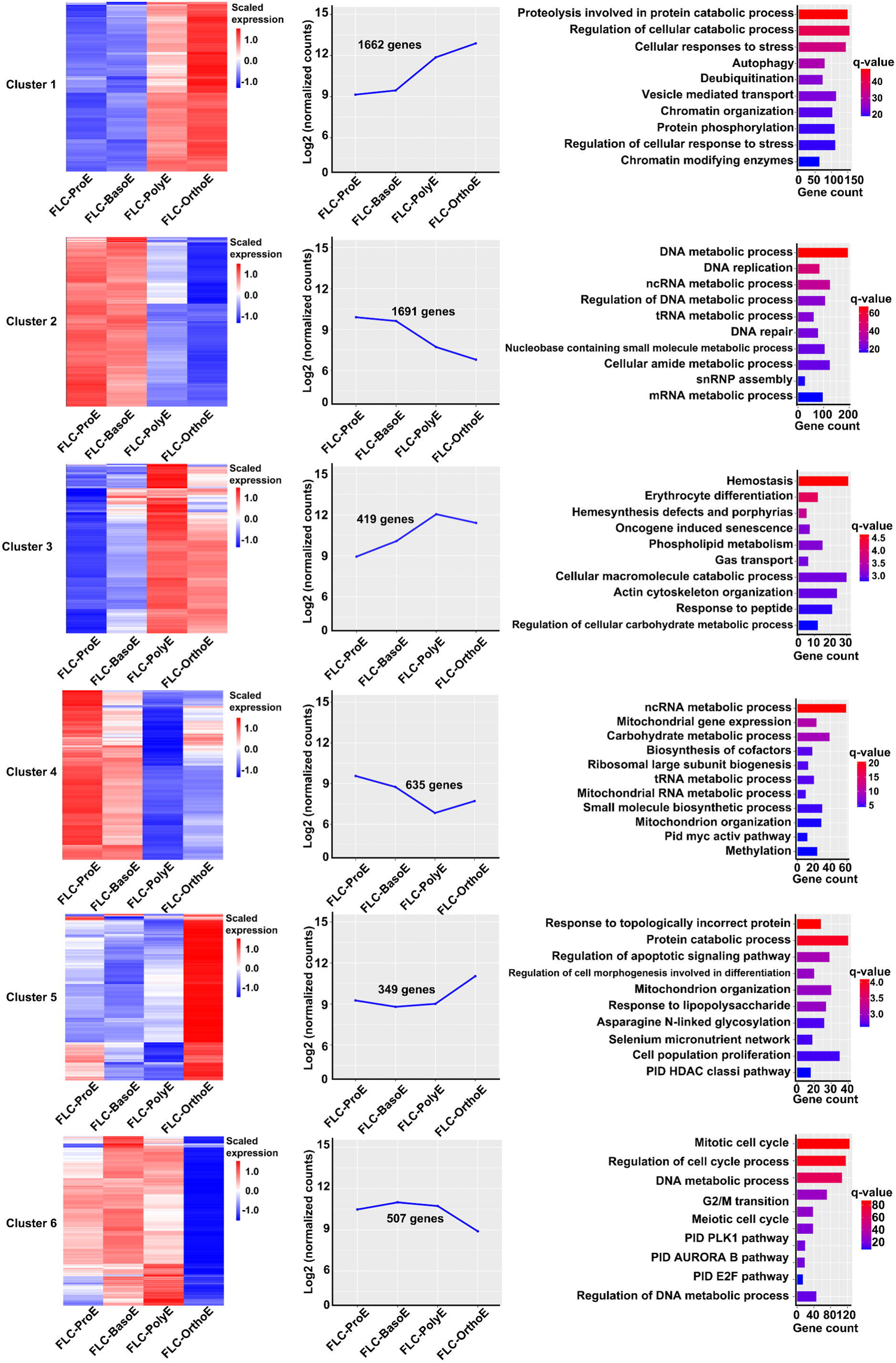
Clusters of differentially expressed genes across stages during *in vitro* terminal erythropoiesis. The gene expression of six identified clusters of all DEGs from adjacent stage comparison in cultured terminal erythropoiesis are shown in a heat map (left panel) and curve plot (middle panel). Enriched GO and pathway terms of genes within clusters are shown in a bar plot (right panel). The q-value represents the log-transformed adjusted *P*.

### Increased lipid metabolism in cultured erythroblasts

In the above-described results, we analyzed DEGs between adjacent stages during *in vivo* or *in vitro* terminal erythropoiesis of human FL. One notable limitation of such an approach is that it may fail to detect differences when the expression of a given gene does not change during terminal erythropoiesis. To clarify this issue, we next compared the gene expression in the same developmental stage *in vivo* and *in vitro*. The numbers of DEGs between primary and cultured erythroblasts increased from ProE and OrthoE with DEG numbers of 553, 790, 2614, and 3599 in ProE, BasoE, PolyE, and OrthoE, respectively. These results imply increased effects of the culture system on erythroblasts along the differentiation process (**Figure 6A**). A list of DEGs at each stage is provided in Table S6. We then performed clustering analyses of all 5148 DEGs identified above and identified 10 clusters (Figures 6B and S3). Notably, among all these clusters, only one cluster (cluster 6) exhibited a stable expression pattern during both *in vitro* and *in vivo* terminal erythropoiesis. Interestingly, the expression levels of genes in this cluster were much higher in cultured erythroblasts than in primary erythroblasts at all stages (Figure 6C). These genes included genes involved in metabolism of lipids (Figure 6D). Given that lipid metabolism is reportedly important in energy production of terminal erythropoiesis [28, 29], these findings suggest that cultured erythroblasts may need higher energy supplies than primary erythroblasts.

**Figure 6.**
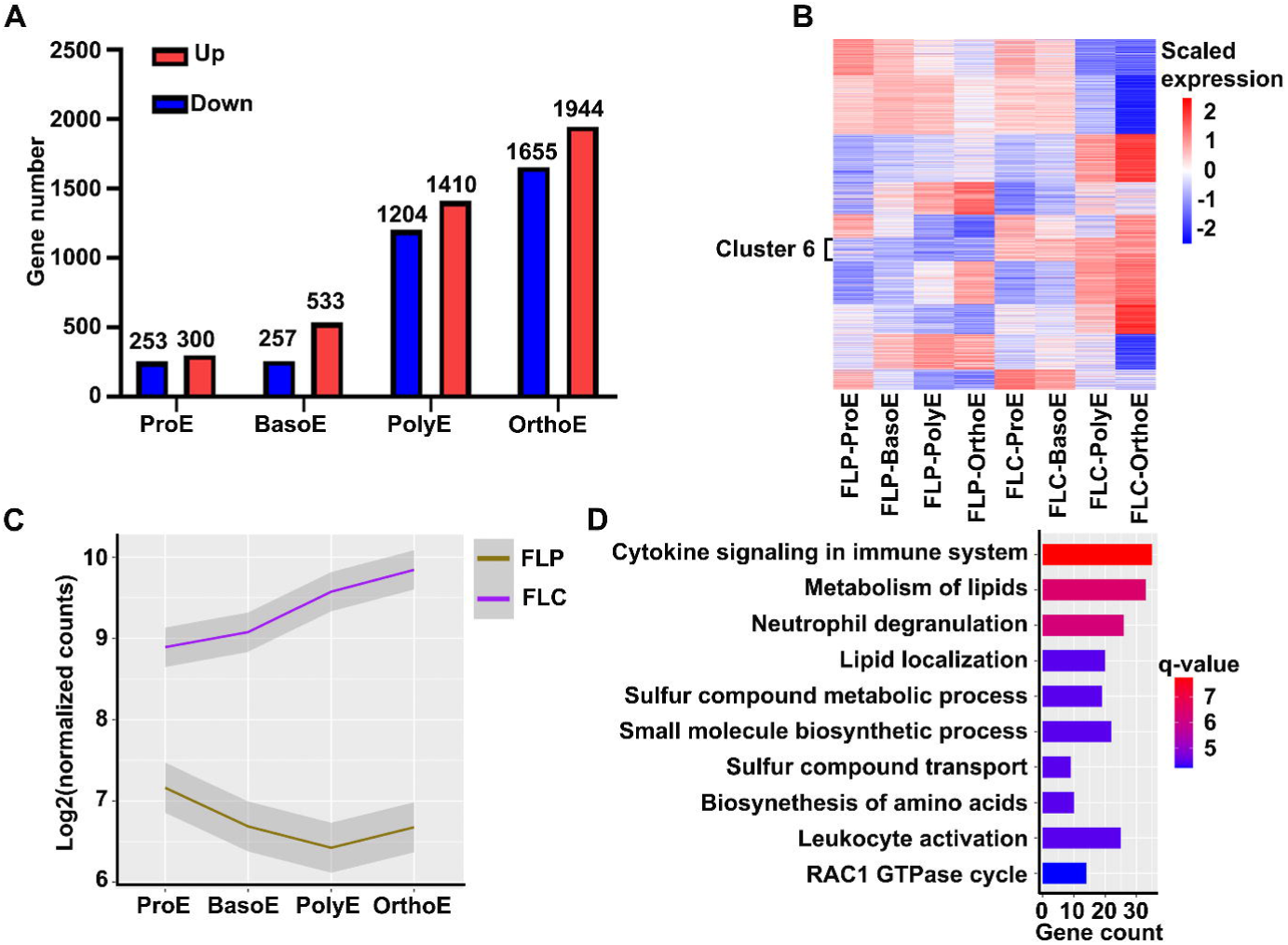
Same-stage comparison between primary erythroblasts and cultured erythroblasts derived from human fetal liver CD34^+^ cells. **A.** Bar plot of DEG numbers of same-stage comparison. **B.** Heat map of gene expression of all DEGs from same stage comparison. **C.** Curve representation of different expression levels of genes in cluster 6. **D.** Bar plot of enriched GO and pathway terms of genes in cluster 6. The q-value represents the log-transformed adjusted *P*.

### Generation and characterization of immortalized erythroid cell line from human FL CD34^+^ cells

In other studies, immortalized erythroid cell lines have been generated from CB, PB, and BM CD34^+^ cells to model neonatal and adult human erythropoiesis [30–32]. However, an immortalized erythroid cell line from human FL is still lacking. To address this, we immortalized FL CD34^+^ cells under erythroid culture conditions utilizing a tetracycline-inducible human papilloma virus (HPV)-E6/E7 system [30–32]. As a control and for comparison purposes, we also immortalized CB CD34^+^ cells under the same conditions. Successful immortalization of the cell line was demonstrated by the fact that the cells continuously proliferated for more than 200 days. **Figure 7**A shows the growth curves of the FL-iEry and CB-iEry with a mean doubling time of 27 and 24 hours, respectively (Figure 7B). Cytospin analyses showed that the immortalized cells morphologically resembled early-stage erythroblasts (Figure 7C). We further examined the expression profile of surface membrane proteins by flow cytometry. As shown in Figure 7D, both FL-iEry and CB-iEry showed moderate expression of GPA; high expression of CD147, CD71, CD36, and α4 integrin; and no expression of band 3. Additionally, neither FL-iEry nor CB-iEry was able to form erythroid colonies (data not shown). Together, these results strongly suggest that the cell lines are immortalized at the ProE stage.

**Figure 7.**
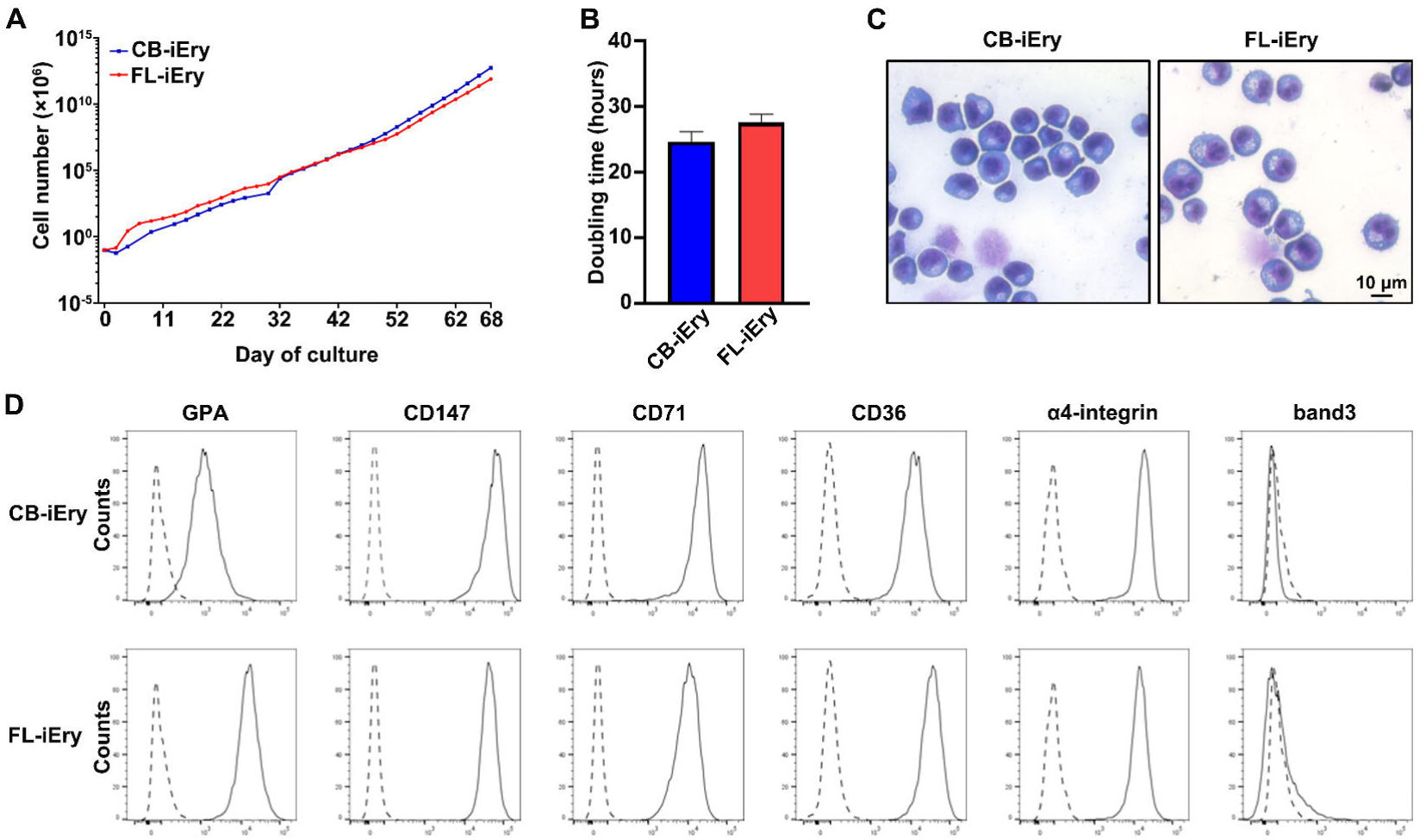
Characterization of erythroid cell lines derived from FL and CB CD34^+^ cells. Growth curves of CB-iEry and FL-iEry. in expansion medium. **B.** Doubling time determined by cell counting. **C.** Representative images of CB-iEry and FL-iEry. Scale bar = 10µm. **D.** Flow cytometric analysis of expression of membrane proteins at cell surface of CB-iEry and FL-iEry. CB-iEry, cord blood CD34^+^ derived immortalized erythroid cell; FL-iEry, fetal liver CD34^+^ derived immortalized erythroid cells.

### Terminal erythroid differentiation profiles of FL-iEry and CB-iEry cell lines

To induce terminal erythroid differentiation, FL-iEry and CB-iEry cells maintained in expansion medium were transferred to differentiation medium. **Figure 8**A shows that in the differentiation medium, the proliferation of FL-iEry and CB-iEry cells lasted for 8 and 6 days, respectively, before they reached plateau. Monitoring the terminal erythroid differentiation process by flow cytometry using band 3 and α4 integrin as markers showed that FL-iEry and CB-iEry underwent terminal erythroid differentiation in a similar manner as demonstrated by the dynamic changes in the expression of band 3 and α4 integrin (Figure 8B). Cytospin analyses showed that morphologically, FL-iEry and CB-iEry progressively matured to BasoE, PolyE, and OrthoE as demonstrated by a decrease in cell size and an increase in chromatin condensation (Figure 8C). Unexpectedly, the OrthoE cells derived from both FL-iEry and CB-iEry failed to enucleate (data not shown). Immortalized human erythroid cell lines provide valuable tools for genetic manipulation to study human erythropoiesis *in vitro* [30–32]. We employed a CRISPR/Cas9 approach to delete two abundantly expressed blood group antigens, CD44 and CD147, in FL-iEry. Gene manipulation was confirmed by genome sequencing (Figure 8D), and deletion of the gene expression was further confirmed by flow cytometry analyses (Figure 8E).

**Figure 8.**
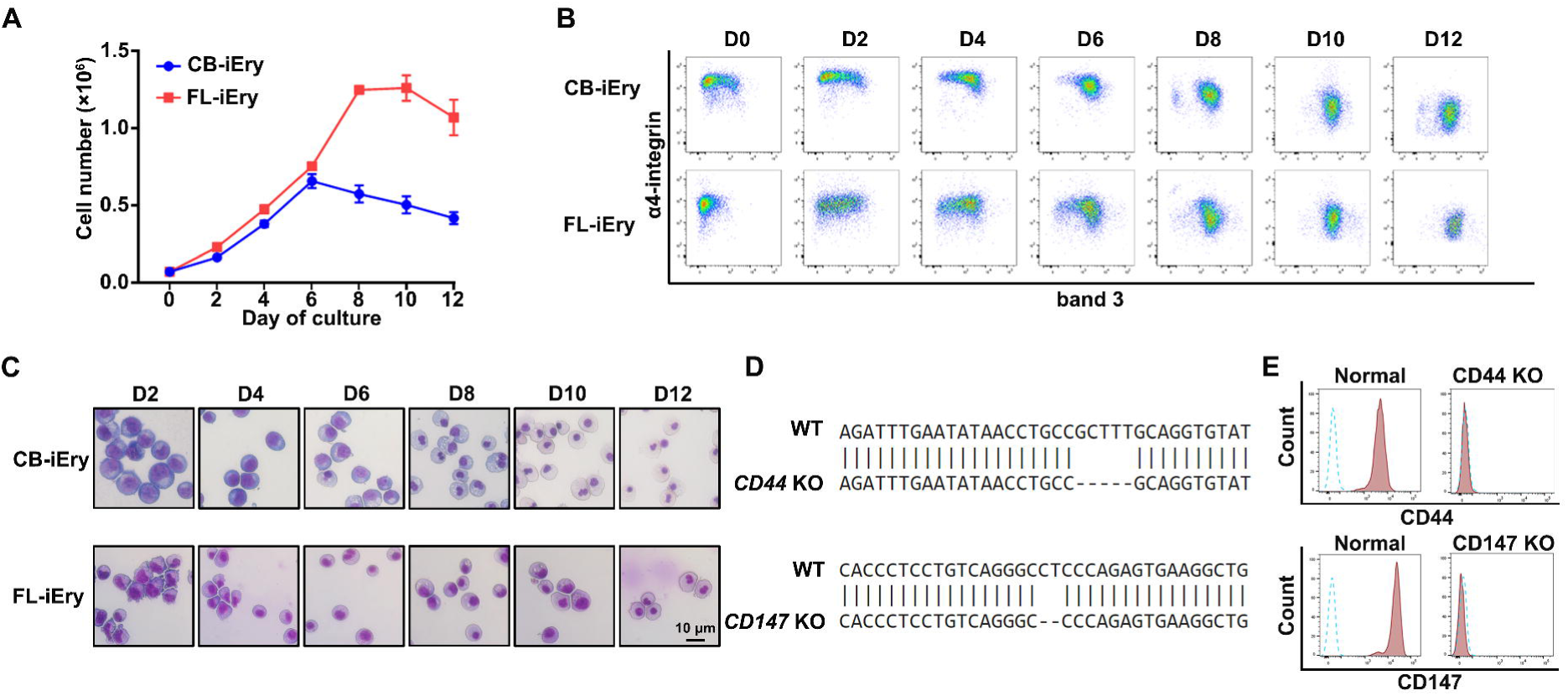
Differentiation and gene knockout in fetal liver and cord blood CD34^+^ derived immortalized erythroid cells. **A.** Growth curves of CB-iEry and FL-iEry. **B.** Representative flow cytometry analysis of band 3 and α4-integrin expression. **C.** Representative cytospin image of erythroblasts from FL-iEry and CB-iEry on different differentiation days. **D.** Genome sequencing in CD44 KO and CD147 KO FL-iEry cell line. **E.** Levels of CD44 and CD147 measured by flow cytometry following knockout of CD44 and CD147 in FL-iEry.

### Transcriptome analyses of immortalized erythroid cell lines revealed immortalization at ProE stage

Next, we conducted RNA-seq analyses of the FL-iEry and CB-iEry cells and compared their transcriptomes with FL CD34^+^ cell-derived and CB CD34^+^ cell-derived erythroblasts at each developmental stage. PCA showed that CB-iEry and FL-iEry were closely clustered and separated from non-immortalized erythroblasts and that CB-iEry and FL-iEry were closest to ProE (Figure S4A). This indicated that CB-iEry and FL-iEry were immortalized at the ProE stage, consistent with the morphology and flow cytometry analyses of the surface markers described above. Although the gene expression profiles of CB-iEry and FL-iEry were closest to ProE, differences are expected to exist between CB-iEry/FL-iEry and ProE. To examine these differences, we compared the transcriptomes of iEry and ProE from the same source: FL-iEry versus FLC-ProE and CB-iEry versus CBC-ProE. In total, 1345 common DEGs were identified, and their expressions are shown in Figure S4B and Table S7. Genes upregulated in iEry were enriched in meiotic synapsis, including genes involved in telomere maintenance such as *TERT* (Figure S4C and D). Given that overexpression of *TERT* leads to the generation of immortalized human cell lines [33], our findings suggest that establishment of the immortalized erythroid cell lines CB-iEry/FL-iEry by the HPV-E6/E7 element occurs at least in part via upregulation of *TERT*.

### Different hemoglobin expression between FL-iEry and CB-iEry

Although FL-iEry and CB-iEry share expression of some common characteristic genes, we were also interested in their differences. PCA showed clear separation between FL-iEry and CB-iEry (**Figure 9**A). In total, 2488 DEGs were identified; 891 genes were upregulated and 1597 genes were downregulated in FL-iEry (Table S8). The DEG expression pattern is represented by a heat map in Figure 9B. GO analyses of the DEGs revealed upregulation of embryonic morphogenesis, reflecting the earlier stage of FL-iEry in development (Figure 9C). Downregulated genes in FL-iEry compared with CB-iEry were enriched in regulation of cell activation and cytokine signaling in the immune system (Figure 9D), implying a possibly lower cytokine response of FL-iEry. Among all the DEGs, we also examined the expression of different hemoglobin genes in Figure 9E. For β-like hemoglobin, adult β hemoglobin *HBB* and *HBD* were significantly downregulated whereas fetal γ hemoglobin *HBG2* and embryonic ε hemoglobin *HBE1* were significantly upregulated in FL-iEry. For α-like hemoglobin, embryonic ζ hemoglobin *HBZ* was significantly upregulated in FL-iEry. The upregulation of embryonic hemoglobins *HBE1* and *HBZ* in FL-iEry compared with CB-iEry further indicates the earlier stage of FL-iEry than CB-iEry in development. Consistent with the upregulation of γ hemoglobin in FL-iEry, the expression of *BCL11A*, which plays an important role in the γ globin to β globin switch [34], was downregulated in FL-iEry compared with CB-iEry. By contrast, the expression levels of *LIN28B* and *IGF2BP1*, both of which promote γ globin expression [35, 36], were increased in FL-iEry compared with CB-iEry. These results indicate that FL-iEry and CB-iEry could be useful cellular models for studying the hemoglobin switch.

**Figure 9.**
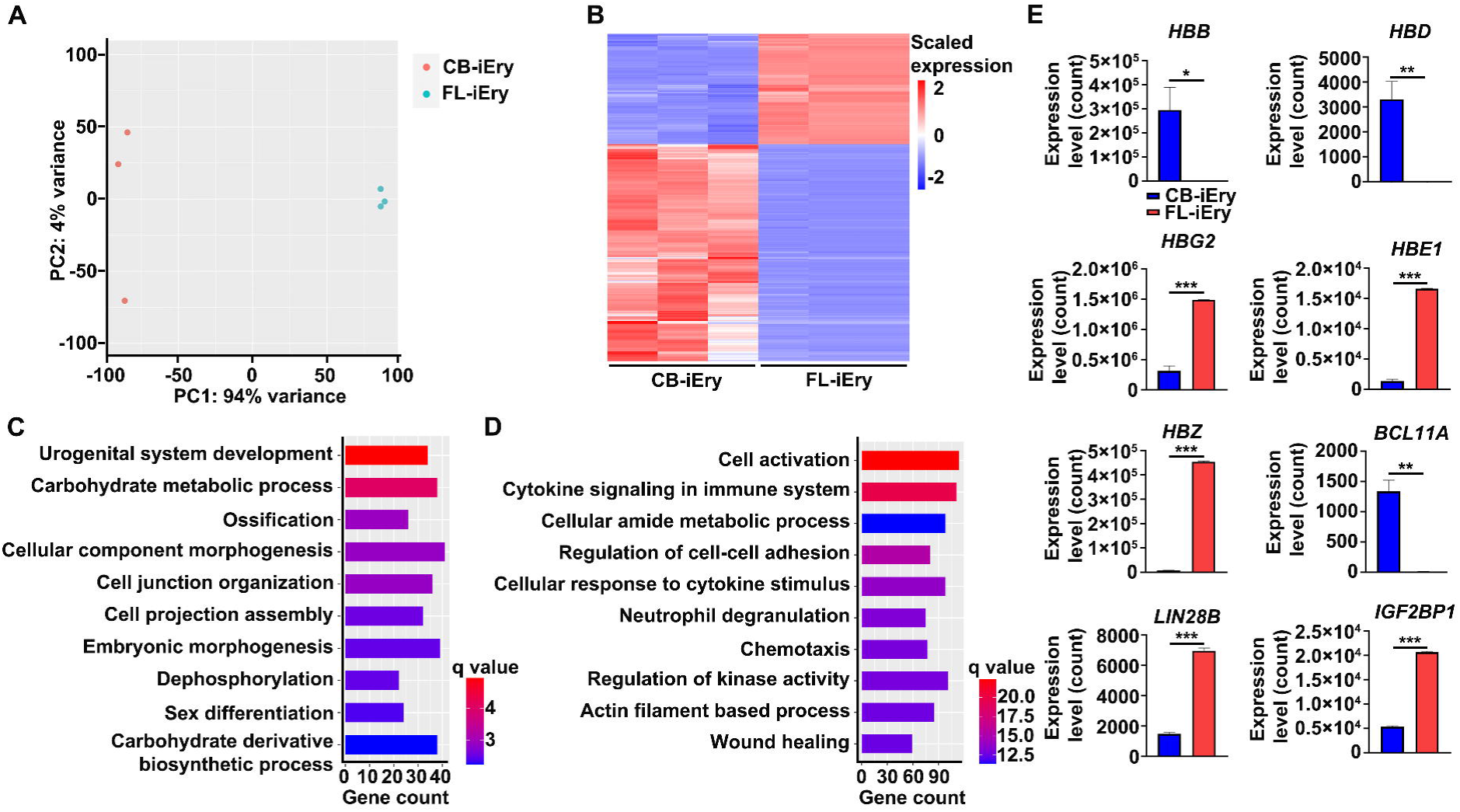
Transcriptome comparison between fetal liver and cord blood CD34^+^ derived immortalized erythroid cells. **A.** Principal component analyses of immortalized cells generated from cord blood and fetal liver CD34^+^ cells. **B.** Heat map of expression of differentially expressed genes between CB-iEry and FL-iEry. **C.** Bar plot of enriched GO and pathway terms of up-regulated genes in FL-iEry. **D.** Bar plot of enriched GO and pathway terms of down-regulated genes in FL-iEry. **E.** Expression of hemoglobin genes and their regulators by normalized counts in CB-iEry and FL-iEry. The q-value represents the log-transformed adjusted *P*.

### Transcriptome comparison of various OrthoE reveals common mechanisms for impaired enucleation

To investigate the mechanisms underlying the inability of OrthoE derived from immortalized erythroid cell lines to enucleate, we performed RNA-seq on OrthoE derived from CB-iEry (CB-iEry-OrthoE) and compared the results with the transcriptomes of other OrthoE, including FL CD34^+^ cell-derived OrthoE (FLC-OrthoE), CB CD34^+^ cell-derived OrthoE (CB-OrthoE) [17], and embryonic stem cell-derived OrthoE (ES-OrthoE) [13]. FLC-OrthoE and CB-OrthoE were able to enucleate, whereas CB-iEry-OrthoE and ES-OrthoE were not. PCA showed clear separation of the four cell groups as well as separation of cells with and without enucleation capability (**Figure 10**A). In total, 1929 DEGs were identified between these two groups (Figure 10B and Table S9). Genes downregulated in cells without enucleation capability were enriched in chromatin organization, histone modification, and mitophagy (Figure 10C). These GO enrichment terms are similar to the terms identified in OrthoE derived from embryonic stem cells that also failed to enucleate [35]. We further examined the expression of genes known to affect enucleation, such as *HDAC5* [26], *FOXO3* [37], *XPO7* [38], *TRIM58* [39], *RIOK3* [40], and *TET3* [20]. The results showed that all of these genes were significantly downregulated in OrthoE without enucleation capability (Figure S5). Together, our findings strongly suggest that defects in chromatin organization, histone modification, and mitophagy are common mechanisms for the inability of immortalized OrthoE and ES-OrthoE to enucleate.

**Figure 10.**
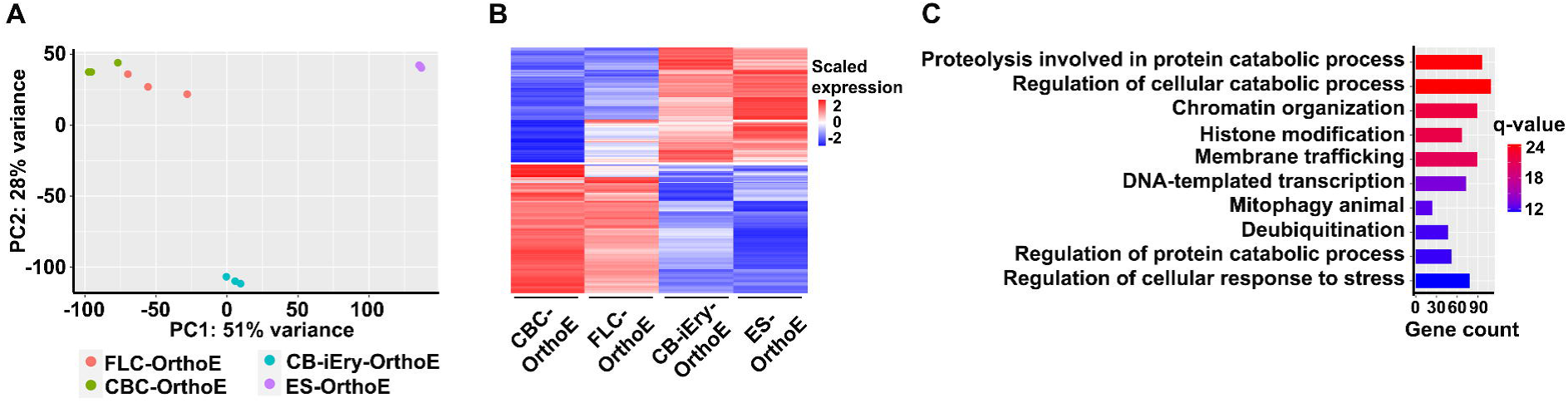
Transcriptome comparison of orthochromatic erythroblasts from different sources. **A.** Principal component analyses of OrthoE from four different sources. **B.** Heat map of expression of common DEGs between cells with and without enucleation capability. **C.** Bar plot of enriched GO and pathway terms of down-regulated genes in cells without enucleation capability groups. The q-value represents the log-transformed adjusted *P*.

## Discussion

The FL is the key erythropoietic organ in the fetus. In contrast to extensive studies on mouse FL erythropoiesis [41, 42], very little is known about human FL erythropoiesis because of limitations in obtaining human FL tissue. In this study, we performed comprehensive characterization and stage-specific transcriptome analyses of human FL terminal erythropoiesis using primary human FL erythroblasts, *in vitro* cultured erythroblasts derived from human FL CD34^+^ cells, and a human FL CD34^+^ cell-derived immortalized erythroid cell line. Our study revealed novel aspects of human FL erythropoiesis. The transcriptomes we generated and the immortalized erythroid cell lines we established provide resources for future studies.

RNA-seq analyses of human erythroid cells have been performed using cultured erythroid cells *in vitro* from CB or PB CD34^+^ cells [17, 43]. Our study provides the first comprehensive analysis of the global transcriptome of primary human FL erythroblasts. The FL cells used in this study were obtained from 16 to 20 weeks of gestation. Given that same markers (Ter119, CD44) can be used to distinguish erythroblasts at different developmental stage from both adult bone and fetal liver [9, 44], we speculate that these markers probably will not change that much during fetal development. However, the transcriptomic profiles will likely to change. Unfortunately, we do not have samples available to perform some tests. Comparison of the transcriptomes between the primary human FL erythroblasts and the *in vitro* cultured erythroblasts derived from human FL CD34^+^ cells revealed several conserved features between *in vivo* and *in vitro* human FL terminal erythropoiesis. First, the numbers of expressed genes progressively decreased as terminal erythropoiesis proceeded. Decreasing numbers of expressed genes were also revealed in our previous RNA-seq analyses of erythroid cells derived from CB CD34^+^ cells [17], indicating a decrease in transcription activity during terminal erythropoiesis. Other commonly changed pathways include upregulation of autophagy and chromatin modification, suggesting that these conserved biological processes are the core mechanisms for terminal erythropoiesis. In addition to conserved biological processes, we also found differences between *in vivo* and *in vitro* terminal erythropoiesis. First, the overall trajectory of *in vivo* and *in vitro* terminal erythroid differentiation was different. Whereas *in vivo* erythropoiesis showed a continuous differentiation process, *in vitro* erythropoiesis showed a two-part separation of the process (ProE/BasoE as one part and PolyE/OrthoE as the other part). Second, while the most significant changes in gene expression were detected between ProE and BasoE *in vivo*, the most significant changes were seen between BasoE and PolyE *in vitro*. Third, more DEGs were detected during *in vitro* than *in vivo* terminal erythropoiesis, particularly in the PolyE and OrthoE stages. One dramatic difference is that the pathway involved in lipid metabolism was significantly upregulated in cultured erythroblasts at all stages. Together, our results provide strong suggestions regarding the environmental effects on gene expression during terminal erythropoiesis.

Immortalized erythroid cell lines have been established from CB, PB, and BM CD34^+^ cells [30–32]. However, the mechanisms by and stages at which the cell lines are immortalized remain unclear. Our cytospin analyses, flow cytometry analyses of surface proteins, and global transcriptomic analyses of immortalized FL-iEry and CB-iEry demonstrated that the cells were immortalized at the ProE stage, indicating their usefulness for the study of terminal erythropoiesis but not for early-stage erythropoiesis. Moreover, the finding that telomere maintenance genes were upregulated in FL-iEry and CB-iEry compared with non-immortalized ProE suggests that establishment of immortalized erythroid cell lines by the HPV-E6/E7 element occurs at least in part via upregulation of *TERT*. Another interesting finding of our study is FL-iEry and CB-iEry expressed different types of hemoglobin and their regulators. The specific expression of hemoglobin *HBZ* in FL-iEry suggests potential application of FL-iEry to studies of the α-like hemoglobin switch.

The enucleation rate of OrthoE derived from ES cells, induced pluripotent stem cells, or immortalized erythroid cells is usually very low [13, 30, 45, 46]. However, the mechanisms underlying the impaired enucleation ability are largely unclear. We compared the transcriptomes of CB-iEry-derived OrthoE, which failed to enucleate, with that of CB CD34^+^ cell-derived OrthoE, which were able to enucleate [20], and found that chromatin organization and mitophagy were significantly downregulated in OrthoE without enucleation capability. Interestingly, downregulation of chromatin organization and mitophagy was also detected in ES-derived erythroblasts that exhibited very low enucleation ability [13]. These findings indicate that defects in chromatin organization and mitophagy are key contributors to the inability of erythroblasts to enucleate. Strategies targeting these pathways should help to improve enucleation.

## Materials and methods

### Antibodies

Mouse monoclonal antibodies to human band 3 and Kell were generated in our laboratory [3, 20, 21]. Commercial antibodies included BV421-conjugated CD45 (563879; BD Biosciences, Franklin Lakes, NJ), PE-conjugated CD235a/GPA (555570; BD Biosciences), APC-conjugated CD235a/GPA (551336; BD Biosciences), PE-Cy7-conjugated CD123 (25-1239-42; Invitrogen, Carlsbad, CA), APC-conjugated CD49d (304307; BioLegend, San Diego, CA), PE-conjugated CD34 (555822; BD Biosciences), APC-conjugated CD36 (550956; BD Biosciences), FITC-conjugated CD36 (555454; BD Biosciences), PE-conjugated CD49e (103805; BioLegend), FITC-conjugated CD47 (556045; BD Biosciences), PE-conjugated CD71 (555537; BD Biosciences), AF647-conjugated CD147 (562551; BD Biosciences), and Hoechst33342 (561908; BD Biosciences).

### Preparation of human FL single-cell suspension

Human FLs of gestational age ranging from 16 to 20 weeks were obtained from Advanced Bioscience Resources, Inc. (Alameda, CA). Single human FL cells were prepared as described previously [47]. Briefly, FL tissues were cut into small pieces, suspended in warm Hank’s solution plus 0.2% collagenase type IV (C5138; Sigma-Aldrich, St. Louis, MO), 2 U/mL DNase I (4716728001; Sigma-Aldrich), 3.26 mM CaCl_2_ (499609; Sigma-Aldrich), and 40 mM HEPES (15630080; Gibco, Grand Island, NY). The tissues were incubated at 37℃ for 30 minutes. The cells were diluted by Iscove’s Modified Dulbecco’s Medium (IMDM) (12440053; Gibco) and passed through a 70-µm cell strainer. Centrifugation was performed at 100 × *g* for 3 minutes. The supernatant was collected, and single cells were obtained.

### Flow cytometry analysis of human FL erythroblasts

Single human FL cells were suspended in staining buffer (phosphate-buffered saline [PBS] / 0.5% bovine serum albumin [BSA] / 2 mM ethylenediaminetetraacetic acid [EDTA]) at 1 × 10^6^ concentration and blocked with 0.4% human AB serum for 10 minutes. The cells were incubated with BV421-conjugated CD45, FITC-conjugated band 3, PE-conjugated GPA, and APC-conjugated α4-integrin at room temperature in the dark. After 15 minutes, the cells were washed with staining buffer by centrifugation at 300 × *g* for 5 minutes. The supernatant was discarded, and the cell pellet was resuspended in staining buffer with 7-AAD (559925; BD Biosciences). The cells were analyzed within 1 hour of staining using a BD LSRFortessa cell analyzer (BD Biosciences).

### FACS of human FL erythroid precursors

Single human FL cells were separated by Ficoll density gradient centrifugation at 400 × *g* for 30 minutes. The mononuclear cells were collected and incubated with CD45 magnetic microbeads (130-045-801; Miltenyi Biotec, Bergisch Gladbach, Germany) for 30 minutes at 4℃ in the dark, according to the manufacturer’s protocol. The CD45^−^ cells were collected using a magnetic-activated cell sorting magnetic beads system. The isolated CD45^−^ cells were suspended in staining buffer (PBS / 2 mM EDTA / 0.5% BSA) and blocked with 0.4% human AB serum for 10 minutes. The cells were stained with PE-GPA, APC-α4-integrin, and FITC-band 3 at room temperature in the dark. After 15 minutes, the cells were washed with staining buffer by centrifugation at 300 × *g* for 5 minutes at 4℃, then resuspended in staining buffer with 7-AAD. The cells were finally sorted on a BD FACSAria fusion cell sorter (BD Biosciences).

### Purification and erythroid culture of human FL CD34^+^ cells

The FL single cells were separated on a Ficoll gradient as described above. CD34^+^ cells were purified from mononuclear cells using CD34 magnetic microbeads. The purified CD34^+^ cells were cultured in a three-phase liquid culture system [3, 20, 21]. In the first phase (D0–D7), CD34^+^ cells were cultured in IMDM with 3% human AB serum,10 μg/mL insulin, 2% human plasma, 200 μg/mL holo-transferrin, 3 IU/mL heparin, 1 ng/mL IL-3, 3 IU/mL erythropoietin (EPO), 10 ng/mL stem cell factor (SCF), and 1% penicillin/streptomycin. In the second phase (D7–D11), the EPO concentration was decreased to 1 IU/mL and the IL-3 was removed. In the third phase (D11–D17), the EPO concentration was decreased to 1 IU/mL, the holo-transferrin concentration was increased to 1 mg/mL, and the IL-3 and SCF was removed. The cells were cultured at 37℃ in 5% carbon dioxide.

### Monitoring of *in vitro* erythropoiesis of human FL CD34^+^ cells by flow cytometry analyses

For analysis of the early stage of erythropoiesis, 0.2 × 10^6^ cells in 25 μL staining buffer (PBS / 2 mM EDTA / 0.5% BSA) were stained with FITC-CD36, PE-CD34, APC-GPA, and PE-Cy7-IL3. To monitor terminal erythroid differentiation, 0.2 × 10^6^ cells in 25 μL staining buffer were stained with PE-GPA, APC-α4-integrin, and FITC-band 3. The stained cells were analyzed with the BD LSRFortessa system (BD Biosciences) as described above.

### Erythroid cells colony assay

Colony assay was performed as described previously [8, 21]. Briefly, the cells were diluted at a density of 200 cells by StemSpan™ Serum-Free Expansion Medium (SFEM) (Vancouver, BC, Canada) in 1 mL MethoCult H4434 for BFU-E and MethoCult H4330 for CFU-E, and they were then incubated at 37°C in 5% carbon dioxide. CFU-E and BFU-E colonies were counted after 7 and 14 days, respectively.

### Construction of plasmid for immortalization

HPV-E6/E7 genomic DNA was subcloned into a pLVX-Tight-Puro backbone (a gift from Genmedic, China) downstream of the Tet operator by the restriction enzyme cutting site EcoRI (NEB) and BamHI (NEB). A tetracycline-controlled transcriptional activator pLVX-Tet-On Advanced plasmid was also prepared (a gift from Genmedic, China). The plasmids were sequenced by Genewiz sequencing service (Azenta Life Sciences, Burlington, MA).

### Lentivirus preparation, transfection, and immortalization of FL CD34^+^ and CB CD34^+^ cells

Lentivirus was prepared as described in our previous studies [20–22]. For virus transfection, human CD34^+^ cells isolated from FL or CB were cultured in differentiation medium (phase 1 medium of three-phase medium, described above) for 2 days. On day 2, the cells were transduced with lentiviral vector pLVX-Tight-Puro-E6/E7 as described previously. On day 3, the medium was changed to fresh medium. On day 4, transduced cells were selected by puromycin (1 μg/mL) and G418 (400 μg/mL). On day 5, the cells were transferred into expansion medium (SFEM containing 50 ng/mL SCF, 3 IU/mL EPO, 1 μM dexamethasone, and 1 μg/mL doxycycline). The cells were then cultured in the expansion medium for more than 60 days to select out the immortalized cells. The medium was changed to fresh medium every other day, and the cell density was maintained at < 0.2 million/mL. When the growth curve became stable, single cells were sorted into a 96-well plate to select the immortalized cell colonies.

### Terminal differentiation of immortalized erythroid cell lines

For immortalized cell differentiation, the cells were transferred into differentiation medium for 6 days along with doxycycline. After day 6, the cells were cultured in IMDM containing 3% AB serum, 2% PB plasma, 10 μg/mL insulin, 3 IU/mL heparin, 3 IU/mL EPO, 1 mg/mL transferrin, and 1% penicillin/streptomycin.

### CRISPR/Cas9-mediated gene deletion in immortalized cell lines

Single guide RNAs targeting CD44 (forward: CACCGCGTGGAATACACCTGCAAAG, reverse: AAACCTTTGCAGGTGTATTCCACGC) or CD147 (forward: CACCGCGTCAGAACACATCAACGAG, reverse: AAACCTCGTTGATGTGTTCTGACGC) were designed by CRISPR design tools (http://crispr.mit.edu/). Guide RNAs were synthesized by Eurofins Scientific (Luxembourg, UK). Guide RNAs were constructed into a CRISPR-Cas9-V3 vector (kindly provided by Xiangfang Wu at Rockefeller University) at the restriction site of *Bbs*I. Lentivirus packaging and virus transduction procedures were performed as described previously [21, 22]. On day 6 post-transduction, GFP^+^CD44^−^ or GFP^+^BSG^−^ single cells were sorted into 96-well plates. After culturing for 2 weeks, clones were picked up for validation of *CD44* or *BSG* gene knockout by genome DNA sequencing.

### Cytospin and May-Grünwald-Giemsa staining

Cytospins were prepared on coated slides with 0.1 × 10^6^ cells using the Thermo Scientific Shandon Cytospin centrifuge. The slides were then stained with May-Grünwald-Giemsa solution (MG500; Sigma Aldrich) for minutes. After rinsing for 90 seconds in 40 mM Tris buffer at pH 7.4, the slides were stained with Giemsa solution (GS500; Sigma-Aldrich) for 15 minutes. Finally, the slides were rinsed twice with water. The cells were imaged using a Leica DM2000 inverted microscope.

### RNA-seq and analysis

We used a QIAGEN RNA isolation kit (74104; QIAGEN, Dusseldorf, Germany) to extract RNA. Samples with an RNA integrity number of > 9 were used for construction of a complementary DNA library by an Illumina TruSeq kit (Illumina, San Diego, CA). The sequencing strategy was 100 bp paired-end by an Illumina HiSeq 4000. We used the RNA-seq data of CB-derived erythroblasts from a previous study [29]. Low-quality reads were removed and filtered by fastp. Filtered reads were quantified by Kallisto using the hg19 transcriptome index [48]. Pairwise comparison of two groups was performed by DESeq2 [49], and the batch effect was included the design of DESeq2. The cutoffs of a fold change of ≥ 2, adjusted *P* of ≤ 0.05, and ≥ 1 transcript per million at least in one group were adopted to identify DEGs in the pairwise comparisons. Log-transformed normalized counts were used in PCA. GO and pathway enrichment analyses were applied by Metascape [50]. GO terms and pathways with a q-value of < 0.001 were considered significant, and the top 10 terms with the smallest q-values were listed in the results. For transcriptome analysis of primary or cultured terminal erythropoiesis, expression patterns were analyzed on all pairwise adjacent DEGs by the time-course clustering R package TCseq using log-transformed normalized counts. For comparison between primary and cultured terminal erythropoiesis, stage-specific DEGs between the same stage of primary and cultured erythroblasts were identified and clustered by TCseq. For transcriptome analysis, we only used late BasoE as BasoE in the comparison.

## Ethics statement

All experimental procedures involving human cells were approved by Zhengzhou University’s ethics committee (ZZUIRB 2023-184).

## Data availability

The datasets generated in the current study are available in the Genome Sequence Archive at the National Genomics Data Center, Beijing Institute of Genomics, Chinese Academy of Sciences / China National Center for Bioinformatics [51]. The accession number is HRA004488. The data can be accessed through the shared URL: https://ngdc.cncb.ac.cn/gsa-human/s/04VK1h0V.

## CRediT author statement

**Yongshuai Han**: Conceptualization, investigation, resources, writing – original draft. **Shihui Wang**: Conceptualization, formal analysis, investigation, visualization, writing – original draft. **Yaomei Wang**: Conceptualization, investigation, resources. **Yuming Huang**: Conceptualization, investigation, resources. **Chengjie Gao**: Conceptualization, resources. **Xinhua Guo**: Conceptualization, resources. **Lixiang Chen**: writing – review & editing. **Huizhi Zhao**: Conceptualization, software, formal analysis, visualization, writing – original draft. **Xiuli An**: Conceptualization, investigation, writing – original draft, writing – review & editing.

## Acknowledgments

This research was supported by Science and Technology Research Project of Henan (Grant No. 232102311003) and the National Natural Science Foundation of China (Grant No. U1804282). We thank Yujia Jiao for her valuable suggestions and essential help in refining this manuscript. We thank Angela Morben, DVM, ELS, from Liwen Bianji (Edanz) (www.liwenbianji.cn), for editing the English text of a draft of this manuscript.

## Supplementary material

**Figure S1 GPA, band 3, and α4-integrin enable isolation of erythroblasts from human fetal liver**

**A.** Flow cytometry analysis of primary erythroblasts of human fetal liver using GPA, band3, and α4-integrin.

**B.** Representative images of sorted erythroblasts from human fetal liver. Scale bar = 10µm. ProE, Proerythroblasts; BasoE, basophilic erythroblasts; PolyE, polychromatic erythroblasts; OrthoE, orthochromatic erythroblasts.

**Figure S2 PCA of terminal erythroblasts from human fetal liver *in vivo* and *in vitro***

FLP-ProE, fetal liver primary proerythroblasts; FLP-BasoE, fetal liver primary basophilic erythroblasts; FLP-PolyE, fetal liver primary polychromatic erythroblasts; FLP-OrthoE, fetal liver primary orthochromatic erythroblasts; FLC-ProE, fetal liver cultured proerythroblasts; FLC-BasoE, fetal liver cultured basophilic erythroblasts; FLC-PolyE, fetal liver cultured polychromatic erythroblasts; FLC-OrthoE, fetal liver cultured orthochromatic erythroblasts.

**Figure S3 Gene expression and GO enrichment of nine clusters of all differentially expressed genes from same stage comparison between human fetal liver *in vivo* and *in vitro***

Heat map of gene expression of all differentially expressed genes in nine clusters are shown in left panel. Curve representation of different expression levels of genes in nine clusters are shown in middle panel. Bar plot of enriched GO and pathway terms of genes in nine clusters are shown in right panel. The q-value represents the log-transformed adjusted *P*.

**Figure S4 RNA-seq analyses of immortalized erythroid cells**

**A.** Principal component analyses of immortalized erythroid cells and each stage of terminal erythroblasts cultured from cord blood and fetal liver CD34^+^ cells. **B.** Heat map of common differentially expressed genes of two separate comparisons between immortalized erythroid cells and their counterpart ProE from the same source. **C.** Bar plot of enriched GO and pathway terms of up-regulated genes in immortalized cells. **D.** Bar plot of *TERT* expression in ProEs and immortalized erythroblasts by normalized counts. ProE, proerythroblast. The q-value represents the log-transformed adjusted *P*.

**Figure S5 Gene expression comparison of known enucleation related genes in cultured OrthoEs**

Gene expression levels are represented by normalized counts. FLC-OrthoE, fetal liver cultured orthochromatic erythroblasts. CBC-OrthoE, cord blood CD34^+^ cell-derived OrthoE; FLC-OrthoE, fetal liver CD34^+^ cell-derived OrthoE; CB-iEry-OrthoE, cord blood immortalized erythroid cell-derived OrthoE; ES-OrthoE, embryonic stem cell-derived OrthoE; ***, adjusted *P* < 0.0001.

**Table S1 Expressed genes in FL *in vivo* terminal erythropoiesis**

**Table S2 DEGs in FL *in vivo* terminal erythropoiesis**

**Table S3 Expressed genes in FL *in vitro* terminal erythropoiesis**

**Table S4 DEGs in FL *in vitro* terminal erythropoiesis**

**Table S5 Stage-specific upregulated genes in FL *in vitro* and *in vivo* terminal erythropoiesis**

**Table S6 DEGs of same-stage erythroblasts between FL *in vitro* and *in vivo* terminal erythropoiesis**

**Table S7 DEGs between immortalized erythroblasts and cultured proerythroblasts**

**Table S8 DEGs between immortalized erythroblasts derived from CB CD34^+^ and FL CD34^+^ cells**

**Table S9 DEGs between orthochromatic erythroblasts with and without enucleation capability**

## References

[1] Palis J, Robertson S, Kennedy M, Wall C, Keller G. Development of erythroid and myeloid progenitors in the yolk sac and embryo proper of the mouse. Development 1999;126:5073–84.

[2] Chen L, Wang J, Liu J, Wang H, Hillyer CD, Blanc L, et al. Dynamic changes in murine erythropoiesis from birth to adulthood: implications for the study of murine models of anemia. Blood Adv 2021;5:16–25.

[3] Hu J, Liu J, Xue F, Halverson G, Reid M, Guo A, et al. Isolation and functional characterization of human erythroblasts at distinct stages: implications for understanding of normal and disordered erythropoiesis in vivo. Blood 2013;121:3246–53.

[4] Nandakumar SK, Ulirsch JC, Sankaran VG. Advances in understanding erythropoiesis: evolving perspectives. Br J Haematol 2016;173:206–18.

[5] Menon V, Ghaffari S. Erythroid enucleation: a gateway into a “bloody” world. Exp Hematol 2021;95:13–22.

[6] Flygare J, Rayon Estrada V, Shin C, Gupta S, Lodish HF. HIF1alpha synergizes with glucocorticoids to promote BFU-E progenitor self-renewal. Blood 2011;117:3435–44.

[7] Liu J, Zhang J, Ginzburg Y, Li H, Xue F, De Franceschi L, et al. Quantitative analysis of murine terminal erythroid differentiation in vivo: novel method to study normal and disordered erythropoiesis. Blood 2013;121:e43–9.

[8] Li J, Hale J, Bhagia P, Xue F, Chen L, Jaffray J, et al. Isolation and transcriptome analyses of human erythroid progenitors: BFU-E and CFU-E. Blood 2014;124:3636–45.

[9] Zhang H, Wang S, Liu D, Gao C, Han Y, Guo X, et al. EpoR-tdTomato-Cre mice enable identification of EpoR expression in subsets of tissue macrophages and hematopoietic cells. Blood 2021;138:1986–97.

[10] Yan H, Ali A, Blanc L, Narla A, Lane JM, Gao E, et al. Comprehensive phenotyping of erythropoiesis in human bone marrow: Evaluation of normal and ineffective erythropoiesis. Am J Hematol 2021;96:1064–76.

[11] Caulier AL, Sankaran VG. Molecular and cellular mechanisms that regulate human erythropoiesis. Blood 2022;139:2450–9.

[12] Xu C, He J, Wang H, Zhang Y, Wu J, Zhao L, et al. Single-cell transcriptomic analysis identifies an immune-prone population in erythroid precursors during human ontogenesis. Nat Immunol 2022;23:1109–20.

[13] Wang S, Zhao H, Zhang H, Gao C, Guo X, Chen L, et al. Analyses of erythropoiesis from embryonic stem cell-CD34(+) and cord blood-CD34(+) cells reveal mechanisms for defective expansion and enucleation of embryomic stem cell-erythroid cells. J Cell Mol Med 2022;26:2404–16.

[14] Dulmovits BM, Tang Y, Papoin J, He M, Li J, Yang H, et al. HMGB1-mediated restriction of EPO signaling contributes to anemia of inflammation. Blood 2022;139:3181–93.

[15] Wang B, Wang C, Wan Y, Gao J, Ma Y, Zhang Y, et al. Decoding the pathogenesis of Diamond-Blackfan anemia using single-cell RNA-seq. Cell Discov 2022;8:41.

[16] Yu L, Lemay P, Ludlow A, Guyot MC, Jones M, Mohamed FF, et al. A new murine Rpl5 (uL18) mutation provides a unique model of variably penetrant Diamond-Blackfan anemia. Blood Adv 2021;5:4167–78.

[17] An X, Schulz VP, Li J, Wu K, Liu J, Xue F, et al. Global transcriptome analyses of human and murine terminal erythroid differentiation. Blood 2014;123:3466–77.

[18] Gautier EF, Ducamp S, Leduc M, Salnot V, Guillonneau F, Dussiot M, et al. Comprehensive Proteomic Analysis of Human Erythropoiesis. Cell Rep 2016;16:1470–84.

[19] Schulz VP, Yan H, Lezon-Geyda K, An X, Hale J, Hillyer CD, et al. A Unique Epigenomic Landscape Defines Human Erythropoiesis. Cell Rep 2019;28:2996–3009.e7.

[20] Yan H, Wang Y, Qu X, Li J, Hale J, Huang Y, et al. Distinct roles for TET family proteins in regulating human erythropoiesis. Blood 2017;129:2002–12.

[21] Qu X, Zhang S, Wang S, Wang Y, Li W, Huang Y, et al. TET2 deficiency leads to stem cell factor-dependent clonal expansion of dysfunctional erythroid progenitors. Blood 2018;132:2406–17.

[22] Han X, Zhang J, Peng Y, Peng M, Chen X, Chen H, et al. Unexpected role for p19INK4d in posttranscriptional regulation of GATA1 and modulation of human terminal erythropoiesis. Blood 2017;129:226–37.

[23] Giarratana MC, Kobari L, Lapillonne H, Chalmers D, Kiger L, Cynober T, et al. Ex vivo generation of fully mature human red blood cells from hematopoietic stem cells. Nat Biotechnol 2005;23:69–74.

[24] Neildez-Nguyen TM, Wajcman H, Marden MC, Bensidhoum M, Moncollin V, Giarratana MC, et al. Human erythroid cells produced ex vivo at large scale differentiate into red blood cells in vivo. Nat Biotechnol 2002;20:467–72.

[25] Jin H, Kim HS, Kim S, Kim HO. Erythropoietic potential of CD34+ hematopoietic stem cells from human cord blood and G-CSF-mobilized peripheral blood. Biomed Res Int 2014;2014:435215.

[26] Wang Y, Li W, Schulz VP, Zhao H, Qu X, Qi Q, et al. Impairment of human terminal erythroid differentiation by histone deacetylase 5 deficiency. Blood 2021;138:1615–27.

[27] Lee E, Sivalingam J, Lim ZR, Chia G, Shi LG, Roberts M, et al. Review: In vitro generation of red blood cells for transfusion medicine: Progress, prospects and challenges. Biotechnol Adv 2018;36:2118–28.

[28] Gibson JS, Rees DC. Lipid metabolism in terminal erythropoiesis. Blood 2018;131:2872–4.

[29] Huang NJ, Lin YC, Lin CY, Pishesha N, Lewis CA, Freinkman E, et al. Enhanced phosphocholine metabolism is essential for terminal erythropoiesis. Blood 2018;131:2955–66.

[30] Kurita R, Suda N, Sudo K, Miharada K, Hiroyama T, Miyoshi H, et al. Establishment of immortalized human erythroid progenitor cell lines able to produce enucleated red blood cells. PLoS One 2013;8:e59890.

[31] Trakarnsanga K, Griffiths RE, Wilson MC, Blair A, Satchwell TJ, Meinders M, et al. An immortalized adult human erythroid line facilitates sustainable and scalable generation of functional red cells. Nat Commun 2017;8:14750.

[32] Daniels DE, Ferguson DCJ, Griffiths RE, Trakarnsanga K, Cogan N, MacInnes KA, et al. Reproducible immortalization of erythroblasts from multiple stem cell sources provides approach for sustainable RBC therapeutics. Mol Ther Methods Clin Dev 2021;22:26–39.

[33] Sealey DC, Zheng L, Taboski MA, Cruickshank J, Ikura M, Harrington LA. The N-terminus of hTERT contains a DNA-binding domain and is required for telomerase activity and cellular immortalization. Nucleic Acids Res 2010;38:2019–35.

[34] Liu N, Hargreaves VV, Zhu Q, Kurland JV, Hong J, Kim W, et al. Direct Promoter Repression by BCL11A Controls the Fetal to Adult Hemoglobin Switch. Cell 2018;173:430–42.e17.

[35] Basak A, Munschauer M, Lareau CA, Montbleau KE, Ulirsch JC, Hartigan CR, et al. Control of human hemoglobin switching by LIN28B-mediated regulation of BCL11A translation. Nat Genet 2020;52:138–45.

[36] Chambers CB, Gross J, Pratt K, Guo X, Byrnes C, Lee YT, et al. The mRNA-Binding Protein IGF2BP1 Restores Fetal Hemoglobin in Cultured Erythroid Cells from Patients with β-Hemoglobin Disorders. Mol Ther Methods Clin Dev 2020;17:429–40.

[37] Liang R, Campreciós G, Kou Y, McGrath K, Nowak R, Catherman S, et al. A Systems Approach Identifies Essential FOXO3 Functions at Key Steps of Terminal Erythropoiesis. PLoS Genet 2015;11:e1005526.

[38] Figueroa AA, Fasano JD, Martinez-Morilla S, Venkatesan S, Kupfer G, Hattangadi SM. miR-181a regulates erythroid enucleation via the regulation of Xpo7 expression. Haematologica 2018;103:e341–e4.

[39] Thom CS, Traxler EA, Khandros E, Nickas JM, Zhou OY, Lazarus JE, et al. Trim58 degrades Dynein and regulates terminal erythropoiesis. Dev Cell 2014;30:688–700.

[40] Zhang L, Flygare J, Wong P, Lim B, Lodish HF. miR-191 regulates mouse erythroblast enucleation by down-regulating Riok3 and Mxi1. Genes Dev 2011;25:119–24.

[41] Isern J, Fraser ST, He Z, Baron MH. The fetal liver is a niche for maturation of primitive erythroid cells. Proc Natl Acad Sci U S A 2008;105:6662–7.

[42] Fantin A, Tacconi C, Villa E, Ceccacci E, Denti L, Ruhrberg C. KIT Is Required for Fetal Liver Hematopoiesis. Front Cell Dev Biol 2021;9:648630.

[43] Yan H, Hale J, Jaffray J, Li J, Wang Y, Huang Y, et al. Developmental differences between neonatal and adult human erythropoiesis. Am J Hematol 2018;93:494–503.

[44] Chen K, Liu J, Heck S, Chasis JA, An X, Mohandas N. Resolving the distinct stages in erythroid differentiation based on dynamic changes in membrane protein expression during erythropoiesis. Proc Natl Acad Sci U S A 2009;106:17413–8.

[45] Olivier EN, Qiu C, Velho M, Hirsch RE, Bouhassira EE. Large-scale production of embryonic red blood cells from human embryonic stem cells. Exp Hematol 2006;34:1635–42.

[46] Qiu C, Olivier EN, Velho M, Bouhassira EE. Globin switches in yolk sac-like primitive and fetal-like definitive red blood cells produced from human embryonic stem cells. Blood 2008;111:2400–8.

[47] Pourcher G, Mazurier C, King YY, Giarratana MC, Kobari L, Boehm D, et al. Human fetal liver: an in vitro model of erythropoiesis. Stem Cells Int 2011;2011:405429.

[48] Bray NL, Pimentel H, Melsted P, Pachter L. Near-optimal probabilistic RNA-seq quantification. Nat Biotechnol 2016;34:525–7.

[49] Love MI, Huber W, Anders S. Moderated estimation of fold change and dispersion for RNA-seq data with DESeq2. Genome Biol 2014;15:550.

[50] Zhou Y, Zhou B, Pache L, Chang M, Khodabakhshi AH, Tanaseichuk O, et al. Metascape provides a biologist-oriented resource for the analysis of systems-level datasets. Nat Commun 2019;10:1523.

[51] Chen T, Chen X, Zhang S, Zhu J, Tang B, Wang A, et al. The Genome Sequence Archive Family: Toward Explosive Data Growth and Diverse Data Types. Genomics Proteomics Bioinformatics 2021;19:578–83.

