## Supplementary figures and images for "Comprehensive Characterization and Global Transcriptome Analyses of Human Fetal Liver Terminal Erythropoiesis"

### supplemental figure 1

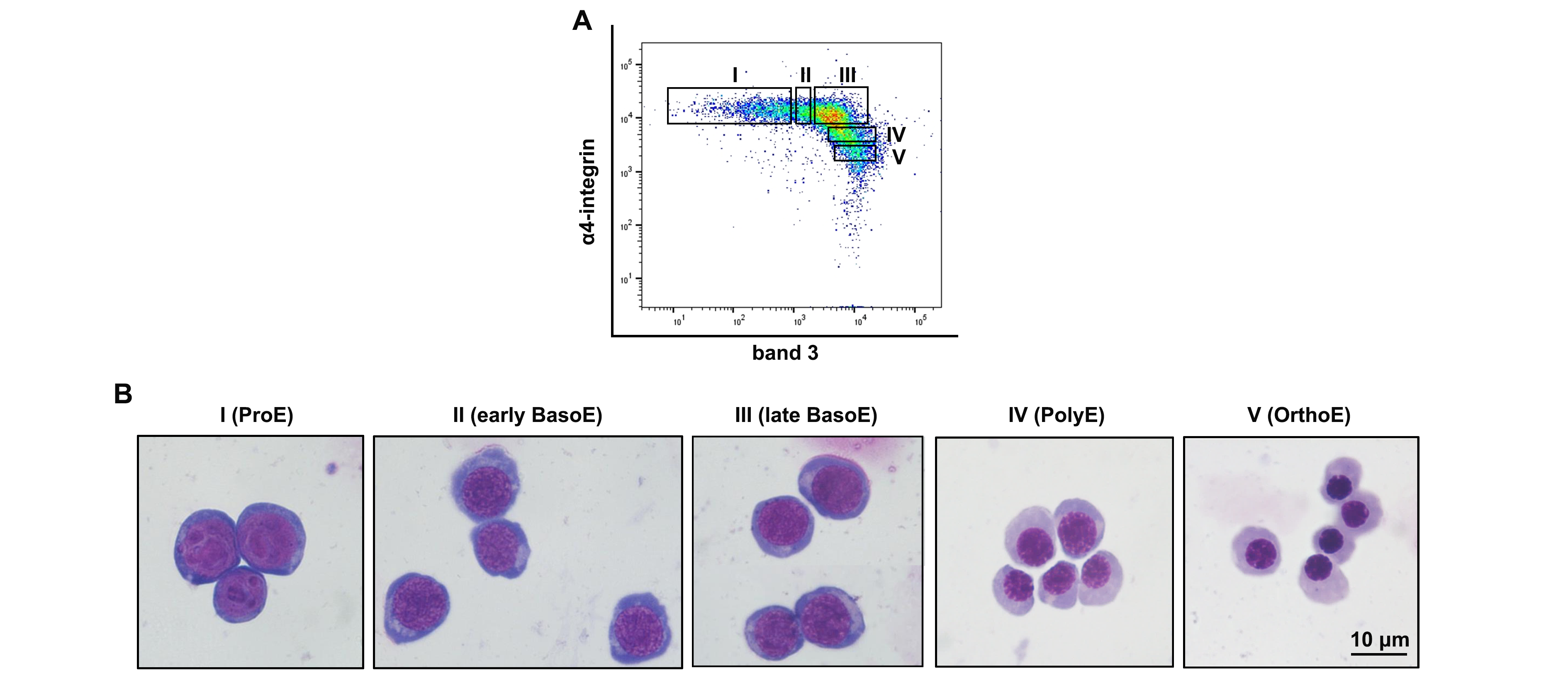

### supplemental figure 2

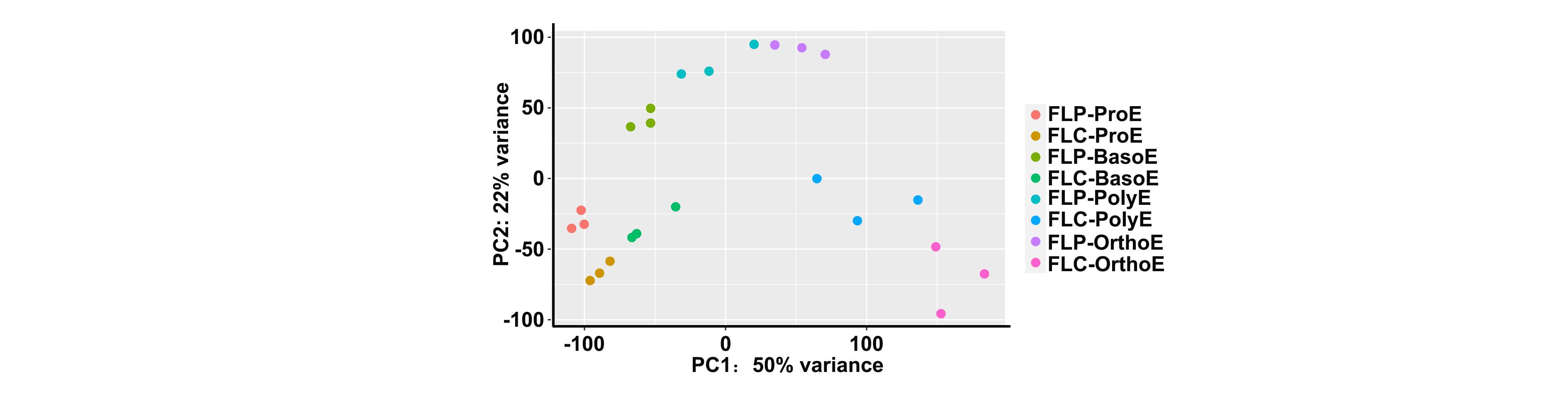

### supplemental figure 3

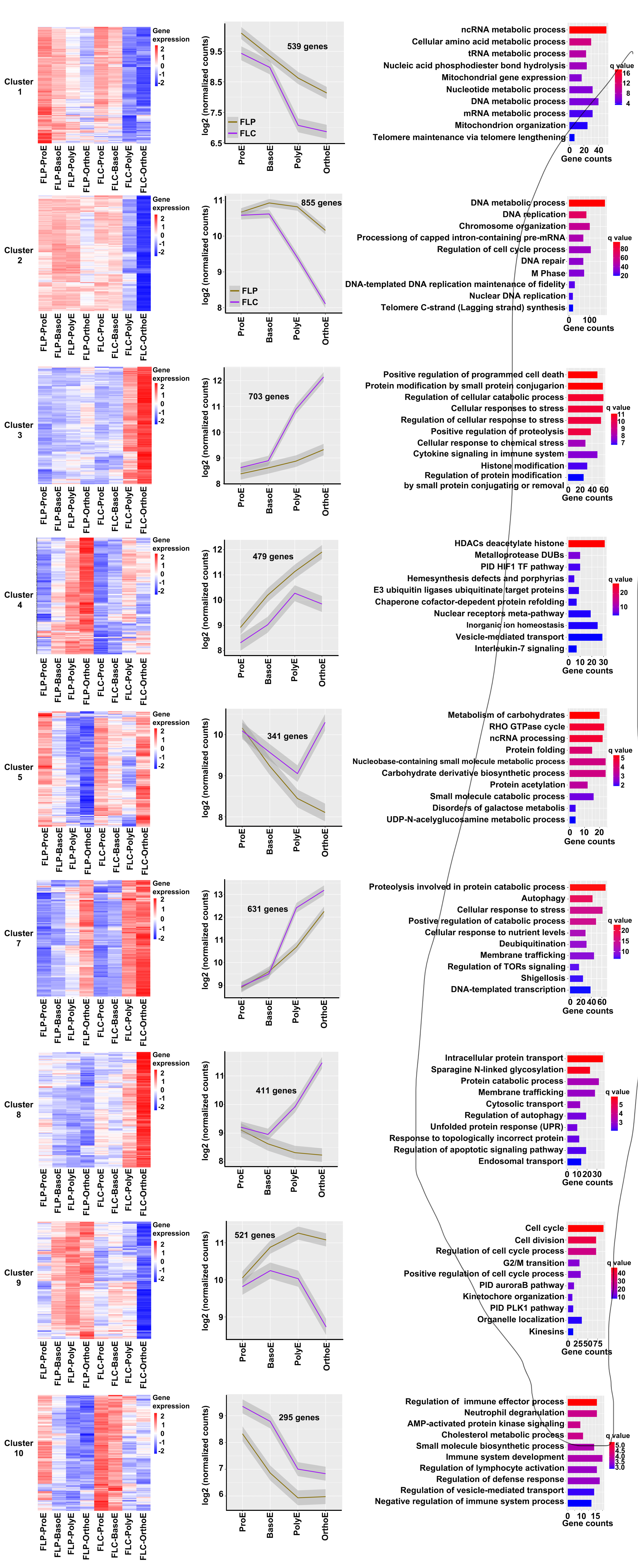

### supplemental figure 4

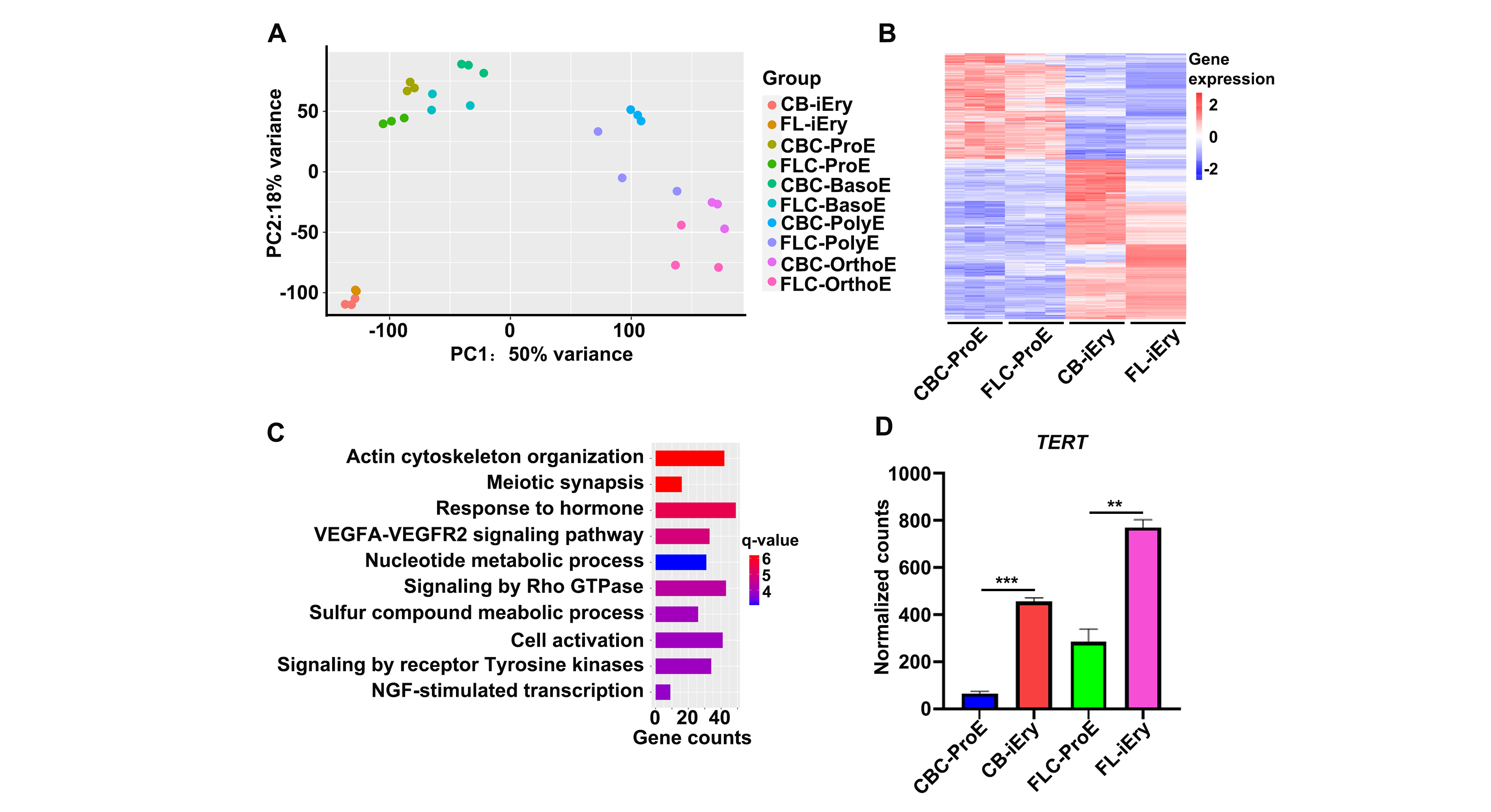

### supplemental figure 5

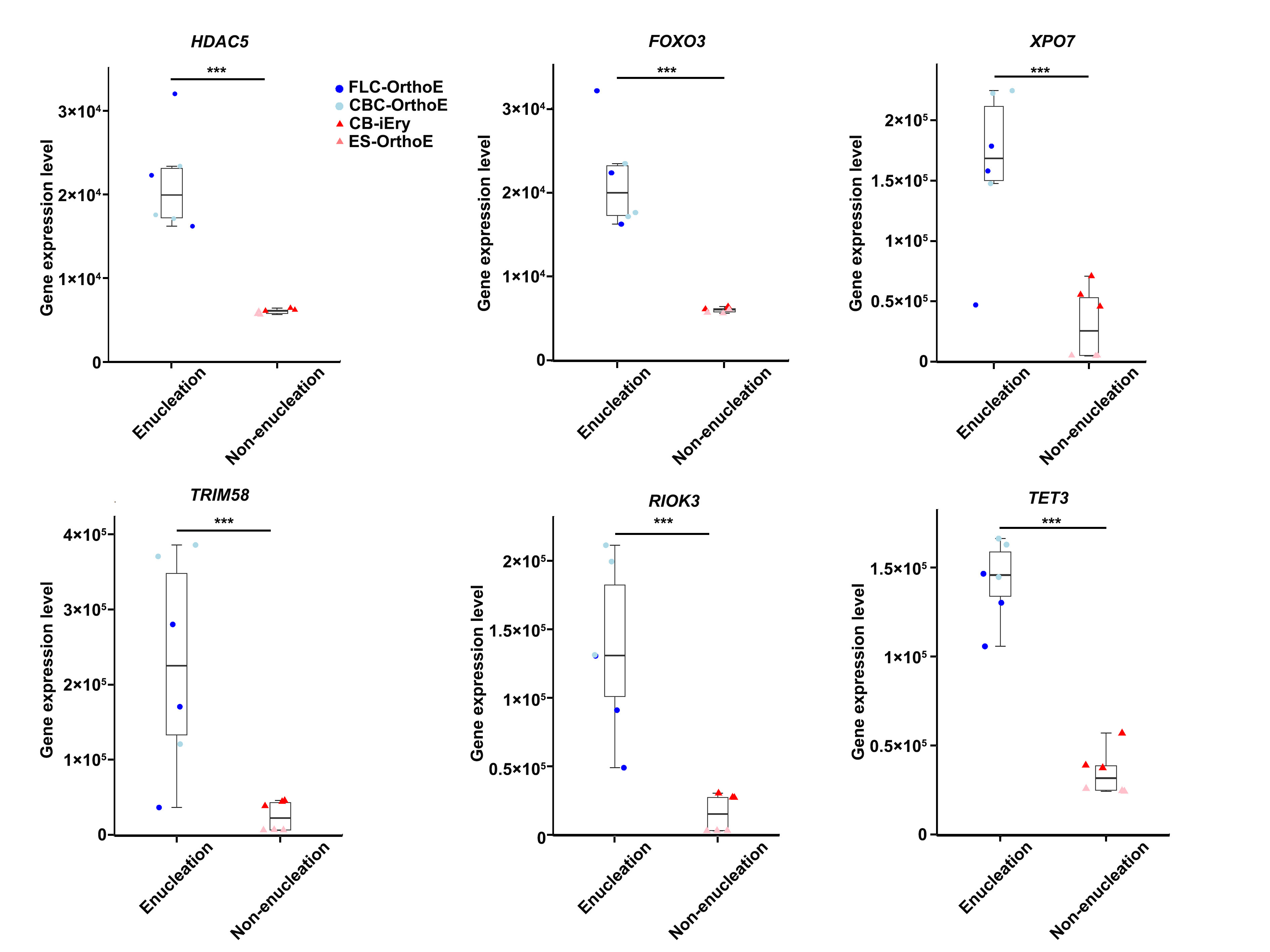
